# Hypermutable hotspot enables the rapid evolution of self/non-self recognition genes in *Dictyostelium*

**DOI:** 10.1101/2025.08.01.668227

**Authors:** Mische Holland, Mayar Ahmed, Janet M. Young, Sean McFadyen, James R. Drurey, Elizabeth A. Ostrowski, Tera C. Levin

## Abstract

Cells require highly polymorphic receptors to perform accurate self/non-self recognition. In the amoeba *Dicytostelium discoideum*, polymorphic TgrB1 & TgrC1 proteins are used to bind sister cells and exclude cheaters, but it remains unknown how cells continually generate this extreme genetic diversity. Here, we created a collection of chromosome-length, whole genome sequences from 10 *Dictyostelium discoideum* isolates and sister species to understand the evolution of the large *tgr* gene family. Our dataset includes AX2-214, a widely used *D. discoideum* lab strain, as well as complete genomes for two *Chlamydia*-like endosymbionts harbored within amoebae. We find that *tgrB1* and *C1* lie in a hypermutational hotspot, with haplotypes that undergo repeated intralocus recombination, duplications, transpositions, and inversions. These structural dynamics are highly localized adjacent to *tgrB* and *C*, resulting in the gain and loss of dozens of genes. The *tgrBC* genes themselves frequently duplicate and recombine, leading to the rapid generation of unique *tgrBC* repertoires. In the broader *tgr* gene family, some genes (e.g. *tgrN*) are single copy and syntenic across all the genomes, whereas others (e.g. *tgrA*) prolifically duplicate at similar rates to *Dictyostelium* transposons. Thus, the *tgr* genes are among the most rapidly evolving families genome-wide. We propose that the intense diversification within the *tgrBC* locus can help explain how these genes acquire such extreme levels of polymorphism, with parallels to the MHC immune genes in mammals and other allorecognition systems. This collection of amoeba genomes is also an ideal dataset for comparative genomics and molecular evolution in Amoebozoa.

**Significance:** The ability to distinguish self from non-self is an essential part of innate immunity and multicellularity. In the amoeba *Dicytostelium discoideum*, cells form transient multicellular structures via aggregation. In the process, they exclude distantly-related cheater strains via highly polymorphic cell surface proteins, TgrB1 and TgrC1. We used this system to ask: how do organisms continually generate new variation in recognition factors? After sequencing a collection of *Dictyostelium spp*. genomes, we found that the *tgrB* and *tgrC* genes lie in an extraordinarily variable locus– a region with such high rates of gene birth and death that genomic similarities are quickly lost, even between closely related isolates. Thus, amoeba cell recognition mirrors self/non-self evolutionary dynamics found across the Tree of Life.

## Introduction

Self/non-self recognition is critical for multicellular coordination. Across diverse organisms, this process has independently evolved to use a variety of recognition proteins as molecular barcodes that can distinguish among cells from closely related individuals(1–5). Mammals use the major histocompatibility complex (MHC) for this purpose(6, 7). In fungi, heterokaryon incompatibility genes mediate cellular fusion barriers(8, 9). Bacterial communities use polymorphic adhesins and toxin-immunity systems to mediate clonal discrimination(10). In all cases, the key features are cell surface proteins that are highly polymorphic and rapidly evolve to generate new lineage identifiers.

*Dictyostelium discoideum* are haploid amoebae that deploy self/non-self discrimination during aggregative multicellular development (Fig. S3A). Transient multicellularity is an asexual process in which individual amoebae aggregate to form fruiting bodies(11), which requires cells to discriminate among potential aggregation partners and prevent exploitation by cheaters(12). This recognition is mediated by a polymorphic transmembrane protein pair, TgrB1 and TgrC1, expressed on the cell surface during aggregation(13–16). Using these “greenbeard” loci, closely related cells with compatible TgrB1/C1 alleles cooperate during development, while incompatible cells are excluded and segregate into distinct fruiting bodies(12, 17–19). This process successfully defends amoebae from neighboring cheater strains(19, 20). Cells that do not express any TgrB1 or C1 proteins are arrested during aggregation and do not complete multicellular development(17).

The *tgr* gene family, which includes *tgrB1* and *tgrC1*, encodes transmembrane proteins with immunoglobulin-like TIG/IPT domains, E-set domains, and Repeats(5). The extracellular TIG domains are critical for cell-cell binding and recognition, while the cytoplasmic domains mediate intracellular signaling(21). Originally classified into 19 informal subfamilies based on sequence similarity and domain architecture(13, 22), only *tgrB1* and *C1* have been functionally characterized. These two genes are extremely polymorphic within *D. discoideum* populations and this sequence diversity correlates with protein binding and developmental compatibility(17, 21). *D. purpureum* has a similarly complex and diverse *tgr* repertoire(23), suggesting that this system of social recognition may be conserved across the genus, but with lineage-specific adaptations.

Notably, genes involved in multicellular development, such as *tgrBC* genes, are thought to evolve more rapidly than other genes in *Dictyostelium(24–26)*, though the lack of high-quality, genetically diverged genomes has limited investigations into the mechanisms of innovation in *Dictyostelium* allorecognition loci. Many *D. discoideum* strains that have been isolated and sequenced are highly genetically similar. As a result, evolutionary analyses have often combined related loci into broad categories of genes(24, 26), reducing both statistical power and explanatory value. Outside the *D. discoideum* AX4 reference genome, high-quality genomic resources have also been limited (Fig. S1). *Dictyostelium* genomes are challenging to assemble due to their high repeat content and low GC content. Although 11 *Dictyostelium* genomes have been generated to date(27), only the assemblies for *D. discoideum* AX4 and *D. firmibasis(28)* are high-quality, near-complete genomes suitable for comparative analyses (Table 1). In addition, because the original AX4 reference strain carries a large chromosome 2 duplication that is absent from other lab strains(29), many researchers have shifted to AX2 lab strains that lack associated genome sequences.

To address these gaps, we generated ten high-quality *Dictyostelium spp*. genomes using long-read sequencing, including both AX2 and species selected to improve phylogenetic sampling between *D. discoideum* and the more distantly related *D. firmibasis(30)*. In the process, we recovered complete genomes of *Chlamydial* endosymbionts from several *Dictyostelium* isolates. When we investigated the evolution of the *tgr* genes, we observed rampant gene turnover and recombination within the *tgrBC* locus, suggesting that these processes drive the generation of new allorecognition alleles. Our analyses also found that certain subfamilies of *tgr* genes are among the most rapidly expanding and diverse genes in the genus, with duplication rates nearing those of the most prominent *Dictyostelium* transposons. In contrast, other *tgr*s are relatively evolutionarily static, suggesting different mutational or selective dynamics within the gene family. These genomic resources for the *Dictyostelium* and Amoebozoa research communities reveal the extensive structural dynamism of allorecognition loci outside of animals, providing a fascinating example of how genetic turnover and local recombination generate and maintain social diversity.

## Results

### *De novo* whole genome assembly of 10 *Dictyostelium* isolates and their endosymbionts

We generated genomes for nine wild *Dictyostelium* isolates collected from across the world as well as the common, axenic *D. discoiduem* lab strain AX2-214 (Table 1). To enable evolutionary comparisons to *D. discoideum* while avoiding re-sequencing genetically similar strains, we prioritized unsequenced group 4 *Dictyostelium* species as well as tropical isolates of *D. discoideum* that were likely cryptic sister species(31). All isolates were sequenced using a combination of Nanopore long reads and Illumina short reads, followed by *de novo* genome assembly (Fig. 1A). In the process, we discovered that three isolates (GS8b, M4B, and PJ11) carried *Chlamydiae* endosymbionts. We obtained complete, circular 1.5 Mbp genomes for two of these endosymbiont isolates, which were similar to recent LF1 and LF3 *Chlamydiae* metagenome assemblies(32) (Fig. S2).

**Table 1.**
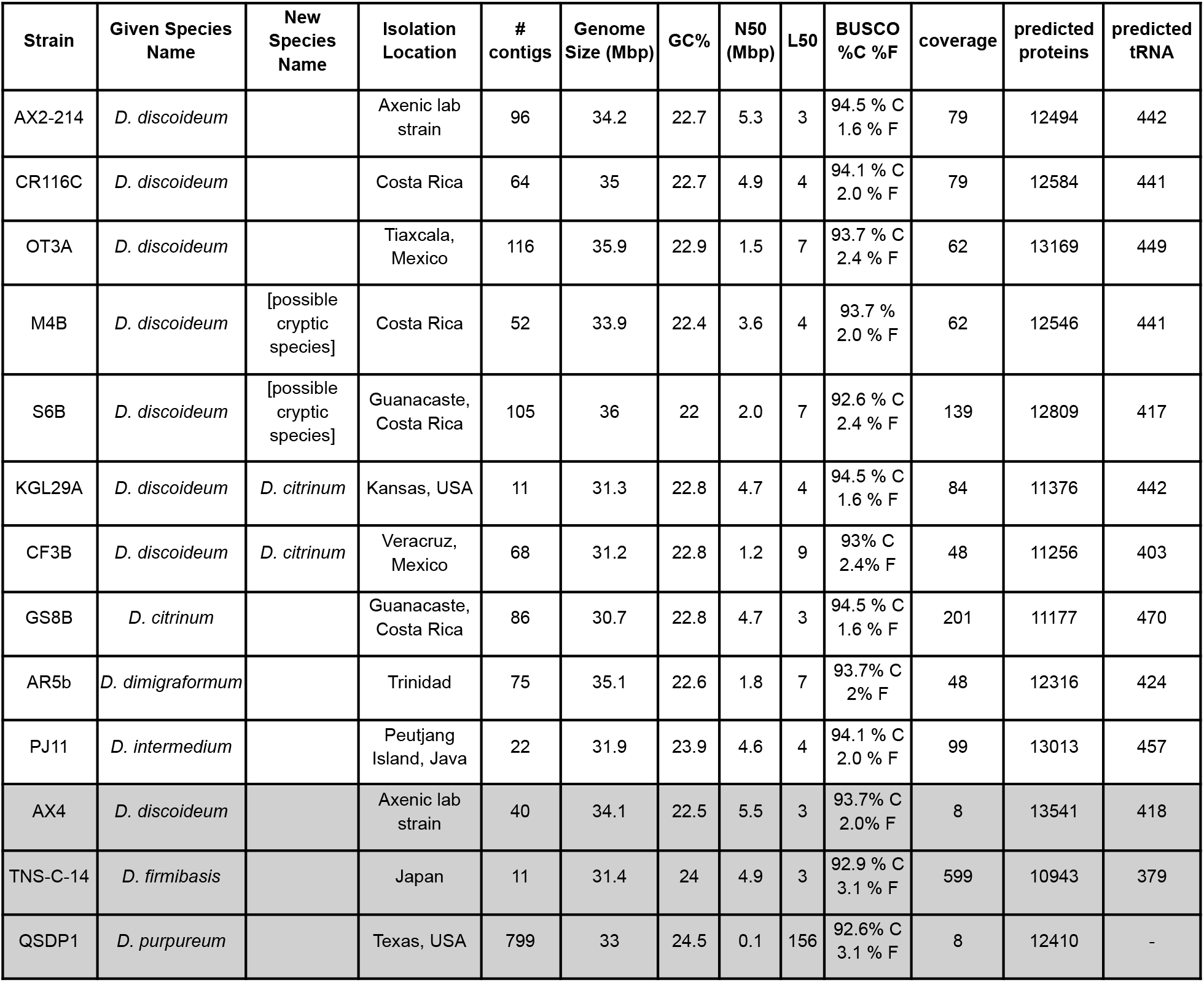
Genome assembly and annotation statistics of *Dictyostelium* genomes. Gray boxes provide statistics for previously published genomes(22, 28, 33). New species names were proposed based on locations on the phylogenetic tree in Figure 1. The AX4 reference genome has six chromosomes. tRNAs were not predicted for *D. purpureum*. BUSCO completeness (%C) and fragmentation (%F) scores are listed for the set of predicted proteins in each genome. See Tables S1 & S3 for more information about sequence quality, rDNA, and mitochondrial DNA.

| Strain | Given Species Name | New Species Name | Isolation Location | # contigs | Genome Size (Mbp) | GC% | N50 (Mbp) | L50 | BUSCO %C %F | coverage | predicted proteins | predicted tRNA |
| --- | --- | --- | --- | --- | --- | --- | --- | --- | --- | --- | --- | --- |
| AX2-214 | *D. discoideum* |  | Axenic lab strain | 96 | 34.2 | 22.7 | 5.3 | 3 | 94.5 % C<br>1.6 % F | 79 | 12494 | 442 |
| CR116C | *D. discoideum* |  | Costa Rica | 64 | 35 | 22.7 | 4.9 | 4 | 94.1 % C<br>2.0 % F | 79 | 12584 | 441 |
| OT3A | *D. discoideum* |  | Tlaxcala, Mexico | 116 | 35.9 | 22.9 | 1.5 | 7 | 93.7 % C<br>2.4 % F | 62 | 13169 | 449 |
| M4B | *D. discoideum* | [possible cryptic species] | Costa Rica | 52 | 33.9 | 22.4 | 3.6 | 4 | 93.7 % C<br>2.0 % F | 62 | 12546 | 441 |
| S6B | *D. discoideum* | [possible cryptic species] | Guanacaste, Costa Rica | 105 | 36 | 22 | 2.0 | 7 | 92.6 % C<br>2.4 % F | 139 | 12809 | 417 |
| KGL29A | *D. discoideum* | *D. citrinum* | Kansas, USA | 11 | 31.3 | 22.8 | 4.7 | 4 | 94.5 % C<br>1.6 % F | 84 | 11376 | 442 |
| CF3B | *D. discoideum* | *D. citrinum* | Veracruz, Mexico | 68 | 31.2 | 22.8 | 1.2 | 9 | 93% C<br>2.4% F | 48 | 11256 | 403 |
| GS8B | *D. citrinum* |  | Guanacaste, Costa Rica | 86 | 30.7 | 22.8 | 4.7 | 3 | 94.5 % C<br>1.6 % F | 201 | 11177 | 470 |
| AR5b | *D. dimigraformum* |  | Trinidad | 75 | 35.1 | 22.6 | 1.8 | 7 | 93.7% C<br>2% F | 48 | 12316 | 424 |
| PJ11 | *D. intermedium* |  | Peutjang Island, Java | 22 | 31.9 | 23.9 | 4.6 | 4 | 94.1 % C<br>2.0 % F | 99 | 13013 | 457 |
| AX4 | *D. discoideum* |  | Axenic lab strain | 40 | 34.1 | 22.5 | 5.5 | 3 | 93.7% C<br>2.0% F | 8 | 13541 | 418 |
| TNS-C-14 | *D. firmibasis* |  | Japan | 11 | 31.4 | 24 | 4.9 | 3 | 92.9 % C<br>3.1 % F | 599 | 10943 | 379 |
| QSDP1 | *D. purpureum* |  | Texas, USA | 799 | 33 | 24.5 | 0.1 | 156 | 92.6% C<br>3.1 % F | 8 | 12410 | - |

**Figure 1.**
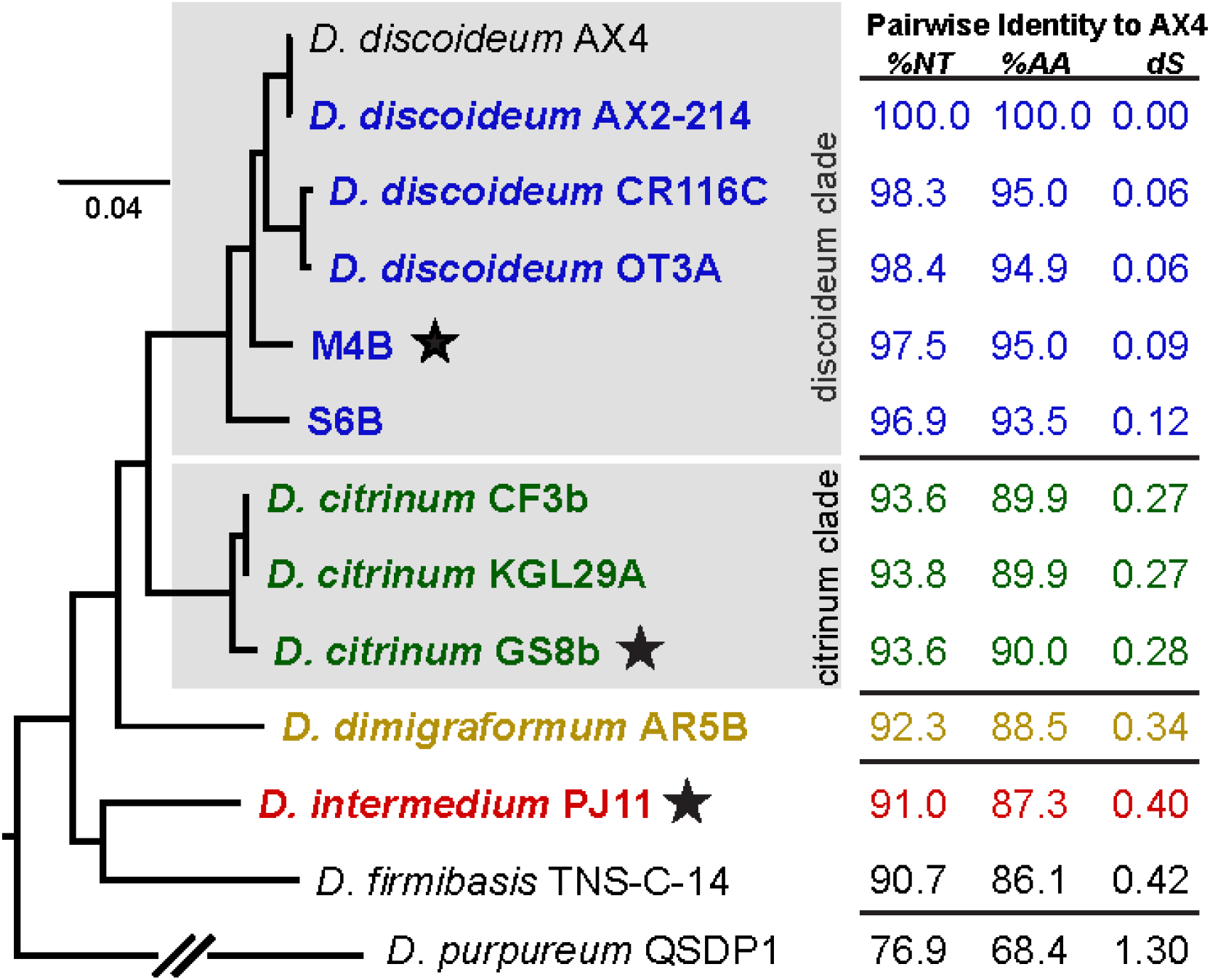
Species tree and diversity statistics for *Dictyostelium* genomes based on 174 BUSCO-identified genes. Genomes assembled in this study are bolded and colored. Pairwise percent identity to the reference AX4 genome and dS comparisons are for BUSCO transcripts (NT) and protein sequences (AA). The presence of *Chlamydia*-like endosymbionts is noted with a black star. All nodes have 100% bootstrap support.

Each *Dictyostelium* isolate had a genome size and GC content roughly similar to the AX4 reference(33), with slightly smaller genomes for the *citrinum*-related strains. Additionally, the new genomes were of similar quality as the original AX4 *D. discoideum* genome, with a similar number of genes and tRNAs (Table 1). The strain divergences (calculated as pairwise nucleotide identity, amino acid identity, or dS relative to AX4) showed that this dataset of 11 genomes is ideal for comparative evolutionary analyses– diverged enough to have acquired genetic diversity, but without reaching sequence saturation (Fig 1). By comparing the pairwise amino acid identities to animals, we estimate that *D. discoideum* AX4 and *D. firmibasis* are approximately as distant as human and mouse(34), between 50-100 million years diverged.

On a phylogenetic tree made from 174 BUSCO conserved genes(35) (Fig. 1), we found that, although CF3b and KGL29A were initially classified as *D. discoideum* (Table 1), they clustered closely with *D. citrinum* GS8b. Additionally, CR116C, OT3A, M4B, and S6B were all genetically diverged from AX4 but clustered together in a *discoideum-*related clade. However, this placement did not reveal whether the amoebae were from the same or different species.

Prior species assignments were based on morphology, which is frequently error-prone for closely related protists. Therefore, we analyzed the mating type loci of each isolate (in the process, discovering some new mating types) and we performed experimental matings among the *discoideum* and *citrinum* clade isolates (Fig. S3). We found that members of the *citrinum* clade formed sexual macrocyst structures with each other, but not with members of the *discoideum* clade (Fig. S3). In addition, AX2 *D. discoideum* formed macrocysts with CR116C and OT3A, but few to none with M4B or S6B. Because of these genetic distances and reductions in mating efficiency, we provisionally call CR116C and OT3A as *D. discoideum*, while M4B and S6B may be cryptic sister species (Table 1).

To evaluate the quality of the genomes, we examined their synteny relative to *D. discoideum* AX4(33) and *D. firmibasis (28)*. For each of the six chromosomes in AX4, most corresponded to only 1 to 3 large, gapless contigs in each genome, indicating that we had generated chromosome-sized assemblies (Fig. 2A). Interestingly, regions corresponding to AX4’s chromosome 2 repeatedly assembled in two separate contigs across all the other genomes, consistent with an AX4-specific fusion and prior findings that lab strains of *D. discoideum* may carry either 6 or 7 chromosomes (36, 37). To determine if our assembled contigs fully extended to the chromosome ends, we searched for markers of *Dictyostelium* centromeres and telomeres. The Dictyostelium Intermediate Repeat Sequence 1 (DIRS1) retrotransposons act as centromeres and are found at the ends of *D. discoideum* chromosomes(33, 38). In the new genomes, many (but not all) contigs contained telocentric DIRS1 clusters and telomeric repeats, suggesting these were chromosome-length (Fig. 2A). The synteny of the genomes also revealed a likely ancestral karyotype for *Dictyostelium* (Fig. 2B), while revealing some lineage-specific chromosomal rearrangements such as chromosome fusions at DIRS1 sites (Fig. 2A, C, & D). Yet, for most genomes, the chromosomal synteny was largely maintained. Using CRAQ(39), we validated that all genomes had high structural assembly quality, and in particular none of the synteny-defined rearrangement breakpoints illustrated in Fig 2 are likely to result from assembly errors (Table S2, Extended Methods). Overall, these analyses suggested that the genome collection had high-quality assemblies that could span complex, repetitive genomic regions.

**Figure 2.**
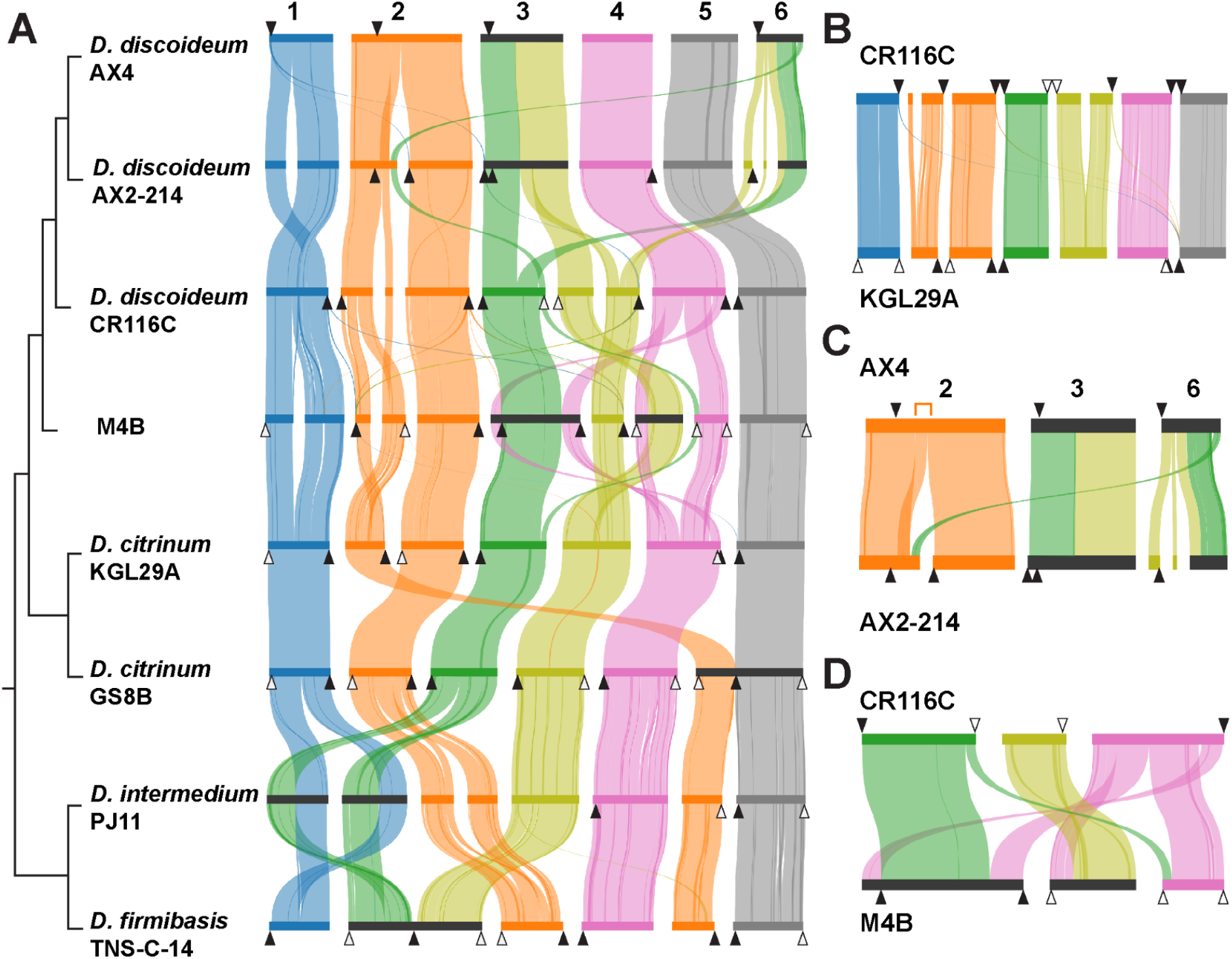
Chromosomal synteny across *Dictyostelium* genomes. **A)** Stacked-parallel synteny plots comparing the best assembled genomes. Every horizontal bar represents a chromosome or contig, colored by ancestral karyotype: chromosome 1 (blue), 2 (orange), 3A (green), 3B (yellow), 4 (pink), and 5 (grey). Ribbons link conserved collinear blocks between genomes, with ribbon width proportional to the number of shared orthologs. Black triangles indicate predicted centromeres (i.e. DIRS1 repeat-rich regions), white triangles indicate predicted telomeres, and horizontal dark gray bars denote inferred chromosomal fusion events. Species tree is provided for reference (not to scale). **B)** Conserved chromosomal synteny between CR116C (*discodium* clade) and KGL29A (*citrinum* clade) suggests that this is the ancestral karyotype. **C)** Compared to the ancestral karyotype and all other genomes, the AX4/AX2 lineage experienced a reciprocal translocation to create chromosomes 3 and 6. Within AX4 chromosome 2, there was also a tandem duplication (indicated with orange bracket) and possible chromosome fusion, as the two halves of chromosome 2 assembled separately in every other genome (see A). **D)** Synteny between *discoideum* clade CR116C and M4B reveals extensive lineage-specific rearrangements involving chromosomes 3A, 3B, and 4 (green, yellow, and pink, respectively).

### tgrs are among the most prolific gene families in Dictyostelium

To understand how genes mediating self/non-self recognition are evolving, we focused on the *tgr* genes, especially *tgrB1* and *C1* that are known to function in allorecognition. In the AX4 genome, the 38 *tgr* genes (and 26 pseudogenes) are divided into 20 subfamilies, named *tgrA, tgrB*, etc., until *tgrS*, as well as a previously unnamed *DDB_G0289561*, which we here name *tgrT1*. Genes within each family are designated with numbers, e.g. *tgrB1, tgrB2*. To identify *tgr* genes across all genomes, we used tblastx to search for sequences similar to AX4 *tgrA-T*, thus identifying 777 homologs that carry the characteristic TIG repeats. After sequence alignment and trimming to remove poorly aligned regions, we found that many of these homologs are truncated or only partially span the alignment length. To remove likely pseudogenes, we filtered out genes that were <50% of the length of the longest homologs, leaving 500 *tgr* genes across the 12 genomes. We then built a phylogenetic tree to categorize the *tgr*s relative to their AX4 homologs (Fig. 3). Ten *tgr* gene families are relatively conserved, found in single copy within each genome, and their clades generally mirror the species tree, suggesting that these *tgrs* are comparatively old and evolutionarily static (*tgrFGLMNPQRS* and *T*, gray clades in Fig 3). In contrast, the *tgrAHI* and *O* genes were numerous, with up to 17 homologs per genome, and these genes frequently inserted in new genomic locations (Fig. S4). While many of these genes are expressed at relatively low levels during clonal development compared to *tgrB1/C1(40, 41)*, at least some of them likely play roles in cell-cell adhesion given their conservation and similar TIG domain architectures.

**Figure 3.**
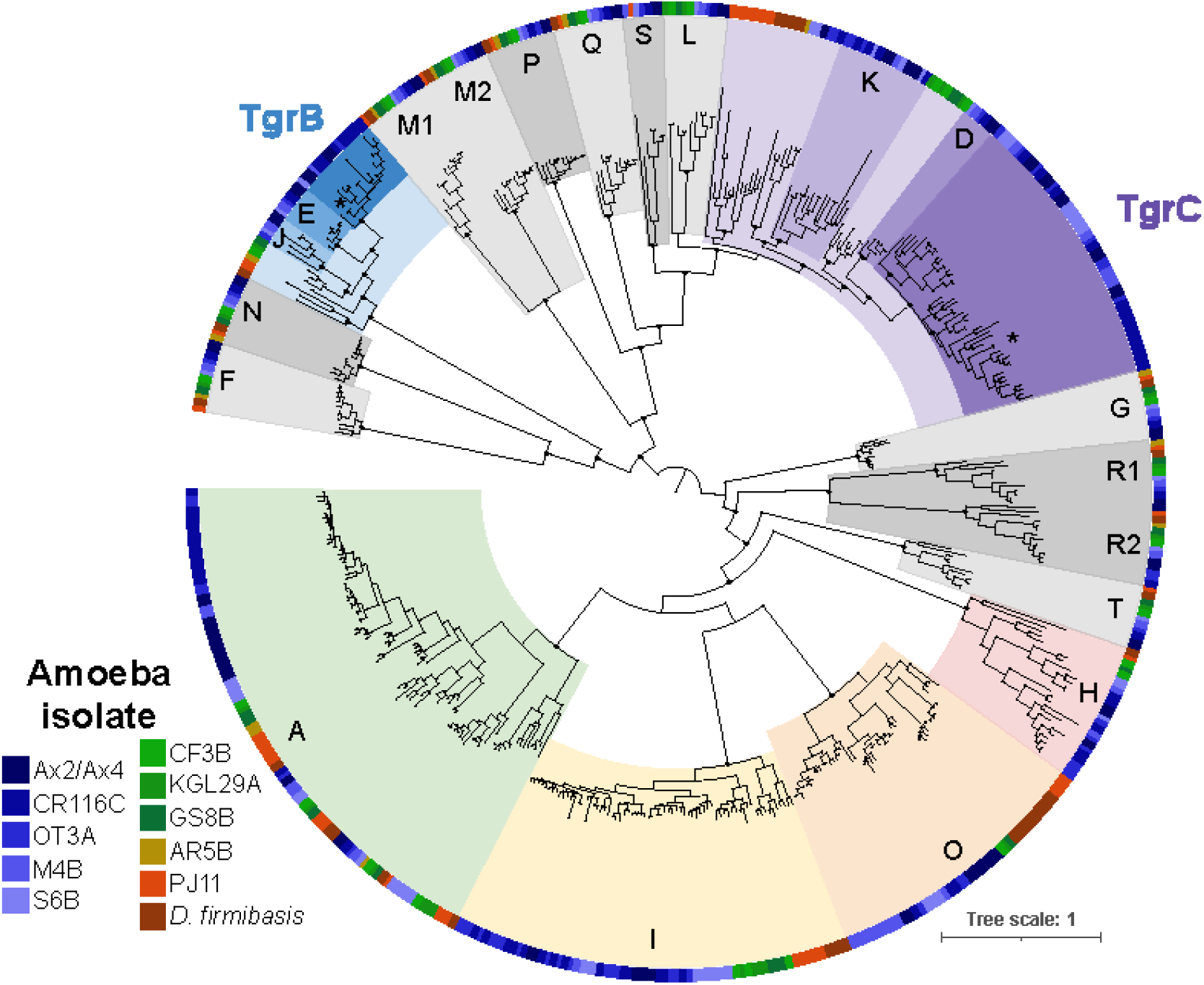
Diversity of Tgrs across 12 *Dictyostelium* spp. Phylogenetic tree of the 500 best-aligning Tgr predicted proteins. Each clade was colored and named according to their relationships to the AX4 homologs. The outer ring is colored according to which genome the Tgr protein came from: blues for *discoideum* clade species, green for *citrinum* clade species, yellow for *D. dimigraformum*, and red/orange for *D. intermedium* and *D. firmibasis*. Gray clades show *tgr* families that are single copy per genome and the clade tree mimics the species tree. Colored clades show rapidly diversifying families: TgrAHIO and TgrBEJ/CDK. The locations of AX4 TgrB1 and C1 are indicated with ^*^. Selected bootstrap supports >90 are shown with black dots for nodes supporting the identity of each clade.

Finally, unlike all other *tgr* families that fell on long, clearly distinct branches, the allorecognition genes *tgrB* and *tgrC* clades were closely related to those of *tgrE/J* and *tgrD/K*, respectively (Fig. 3). When we considered only genes that were one-to-one orthologs with AX4 genes, all of the *tgrB, E, J, C, D*, and *K* genes were restricted to the *discoideum* clade species, suggesting that each of these genes arose from recent gene duplication and diversification. *D. citrinum* and more distantly related species did encode homologs of the *tgrBEJ* and *tgrCDK* families, many of which resided within the *tgrBC* syntenic locus (see below). When we compared our results to *D. purpureum*, we found that *D. purpureum* encoded many lineage-specific *tgr* genes, and only *tgrM* and *tgrN* had clear orthologs in AX4 (Dataset S1, contrary to (23)). Thus, while nearly all *tgr* families have diversified across long timescales, the activities are more intense in *tgrAHIO, tgrBEJ*, and *tgrCDK*.

To evaluate whether the *tgr* gene family expansion is unusual relative to other *Dictyostelium* genes, we categorized all predicted genes into orthology groups. We recovered 10,958 total orthogroups, of which 6,484 contained single-copy genes across all species, representing the *Dictyostelium* core genome (Fig. S5). Consistent with prior studies(33, 42–44), the 15 largest orthogroups include polyketide synthases, actins, and many transposable elements, including the centromere-associated DIRS1 transposon. By comparison, even after dividing the *tgr* genes into subfamilies, the *tgrA* and *tgrCDKLS* are also among the largest orthogroups in *Dictyostelium*, each containing at least 4 predicted genes per genome.

The abundant polymorphisms within *tgrB1* and *tgrC1* are known to be critical to their allorecognition functions (17). To investigate whether other *tgr* genes and *Dictyostelium* isolates show similar patterns of polymorphism, we analyzed the nucleotide and amino acid divergence within the *discoideum* clade genomes. Within the rapidly expanding *tgr* gene families (AHIO, BEJ, and CDK), we observed a broad range of pairwise amino acid identities, from 30% to 80%. This wide range likely reflects the presence of numerous partial gene fragments that may be pseudogenes, similar to the AX4 *tgrB2* and *tgrC2–C5* genes that exhibit low or no expression during development (40, 41) (Fig. S5C). The functions of the TgrAHI and O proteins are unknown, and thus functional orthologs cannot readily be confidently identified.

In contrast, we identified candidate functional loci for the *tgrB* and *tgrC* families by searching for *tgrBC* genes that are: 1) in the same syntenic locus as Ax4 *tgrB1/C1*, and 2) encoded as paired, diverging genes that share a single promoter. Across the 11 isolates, we identified 1-9 such pairs per genome. However, in 13 of the 25 total loci, the *tgrB* genes were shortened or carried large deletions of >50 amino acids. After removing these potential pseudogene loci, we were left with 12 intact loci that we called as *tgrB1/C1*s: two pairs in *D. intermedium* PJ11 and one pair in every other genome. Surprisingly, these curated *tgrB1* genes had a high percent amino acid identity of 80-90% (similar to *tgrFGLMNPQRS* and *T*), whereas this number was 65-70% for their corresponding *tgrC1*s, implying that the evolution of *tgrB1* and *C1* occurs at different rates (Fig. S5C). When we looked at *tgr* diversity within the *discoideum* clade, *tgrC1* had higher nucleotide diversity than other *tgr* genes. Both *tgrB1* and *C1* had a modestly elevated dN/dS relative to other single-copy *tgr* and BUSCO genes, but dN/dS was still <1, making it unclear whether this higher rate of amino acid substitution reflects positive or relaxed selection (Fig. S5F-H). For both *tgrB1* and *C1*, Tajima’s D was slightly greater than 0, but there was not strong evidence of balancing selection. These findings suggest that *tgrB1* and *tgrC1* appear to be evolving at different rates from one another and from the other *tgr* genes, although we did not detect strong signatures of selection.

### Rampant rearrangements in the *tgrBC* locus drive the diversification of allorecognition genes

Given that we did not observe strong signatures of selection, how do *tgrBC* genes generate and maintain their diversity? Unlike other genomic regions, the area surrounding the *tgrB1/C1* pair displayed extremely complex synteny, which made it difficult to understand how individual *tgr* genes were evolving (Fig 2, Fig. S6). Therefore, we leveraged the synteny of flanking marker genes upstream and downstream of *tgrB1* and *tgrC1*. These marker genes, *wacA, chdB*, and *DDB_G0280717*, enclose a region of over 104 kb on AX4 chromosome 3, which we define as the “*tgrBC* locus”. As expected, between AX4 and AX2 lab strains of *D. discoideum*, the *tgrBC* locus was 99.5% identical at the nucleotide level with no rearrangements, and none of the genetic changes were within the coding regions of *tgrB* or *tgrC* genes, demonstrating that we could successfully assemble these complex genomic regions. But in contrast to these two strains, across the other genomes, the *tgrBC* syntenic locus frequently expanded and contracted, ranging from 104 kb to over 250 kb between the marker genes.

The *tgrBC* behaves as a local mutational hotspot, with rampant, intralocus recombination, duplications, inversions, and alterations in gene content within the region (Fig. 4). These rearrangements resulted in unusual, overlapping synteny assignments from the MCScanX software, where the *tgrBC* locus in one genome was syntenic with another genome’s *tgrBC* locus on both the forward and the reverse strand simultaneously, depending on which genes in the locus were considered (Fig. S6). To better understand how these dynamics impacted *tgr* gene diversity, we generated alignments of the *tgrBC* loci between pairs of closely related *Dictyostelium* isolates. Surprisingly, between *D. discoideum* AX4 and OT3A the syntenic genes at the edges of the loci flanked a large, unalignable region downstream of *tgrC1*, wherein both genomes encoded different complements of genes (34kb with 29 genes in AX4 swapped for 138kb with 53 different genes in OT3A, Fig. 4A). These *tgrBC* genomic swaps were common. We found similar unalignable regions downstream of *tgrC1* (and sometimes in additional portions of the *tgrBC* locus) in all other pairwise comparisons across the 11 genomes (Fig. S7). These unalignable regions often contained duplicates or partial gene fragments of *tgrB, tgrC*, and many other neighboring genes in the *tgrBC* locus. When we analyzed pairs of genomes that were more closely related, such as *D. citrinum* KGL/GS8B, the unalignable regions were present, but comparatively small (Fig. 4B). When the genomes were distantly related, as between *D. intermedium* and *D. firmibasis*, the regions were larger and more numerous (Fig. 4C), although these distantly related pairs were more difficult to accurately align given their diverged sequences. Nevertheless, across the *Dictyostelium* genomes, the locus contained novel, jumbled genomic regions adjacent to the paired *tgrBC* genes.

**Figure 4.**
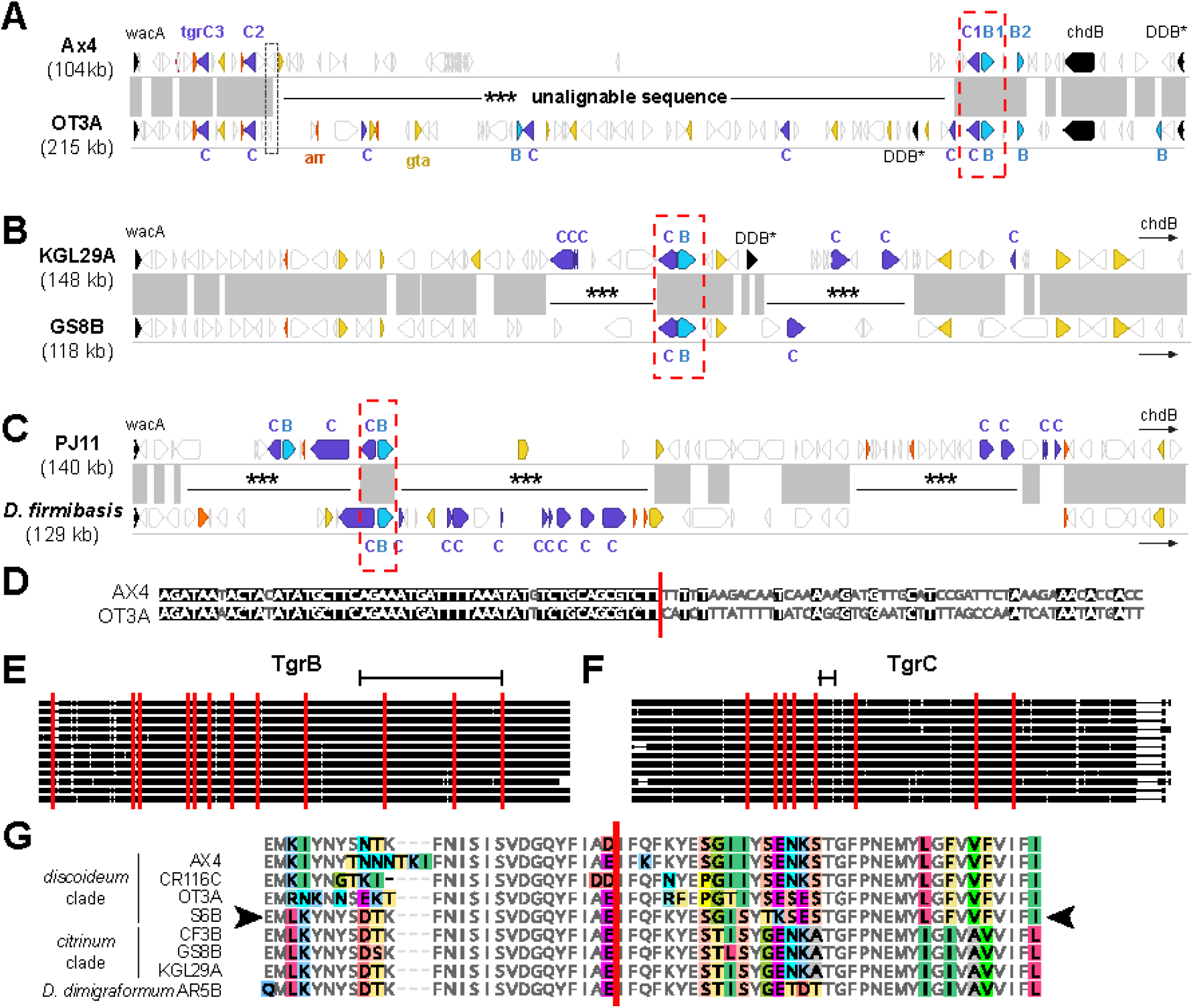
The *tgrBC* locus evolves through recombination and dynamic local gene expansions. **A-C)** Mauve pairwise alignments of *tgrBC* loci, comparing **(A)** AX4 and OT3A in the *discoideum* clade, **(B)** KGL29A and GS8B in the *citrinum* clade, and **(C)** *D. intermedium* PJ11 vs *D. firmibasis*. The length of the region shown is at the left. All genes in the region are illustrated as block arrows (not to scale). Paired, diverged *tgrBEJ/CDK* genes are indicated with red dotted boxes. The marker genes *wacA, chdB*, and *DDB_G0280717* (“DDB^*^”) are in black, with small arrows indicating that the marker gene is downstream of the illustrated region. The *tgrBEJ* (light blue) and *tgrCDK* (purple) genes and gene fragments are colored, as are neighboring genes that are frequently duplicated: *gta* (yellow) and *arr* (orange). Synteny between genomes is shown with the gray boxes, representing regions of at least 30% nucleotide identity. Unalignable regions are indicated with ^***^. **D)** Recombination events at the edges of unalignable regions (red line) are easily detected as transitions between highly similar sequences (left) and regions of divergence. Alignment lies within *DDB_G0280741*, at the unalignable boundary near *tgrC2*, in the black dotted box in panel A. **E & F)** Alignments of curated, intact *tgrB* and *C* sequences identify statistically significant recombination events (vertical red lines), including some within the regions of TgrB/TgrC experimentally determined to mediate recognition (black brackets). Gaps in the alignment are shown as thin horizontal lines while aligned sequences are thick lines. **G)** Example TgrB recombination site (red line), where the S6B sequence (arrowheads) matches the lower citrinum clade sequences on the left, but matches the top three discoideum clade sequences on the right.

Genome rearrangements can result from multiple types of regulated or pathological genome instability. Because these large genomic regions appeared in a short period of time, we tested whether the unalignable regions were moving into the tgrBC locus from another genomic site by using blastn to search for the unalignable sequences elsewhere in the genome. However, other than the *tgrBC* locus itself, none of the resulting hits were larger than a single gene, suggesting that these extended regions did not originate outside the locus. We also found no evidence of specific DNA motifs associated with the unalignable regions, and the boundaries of the regions were found both within and between genes (e.g,. Fig 4D). Two genomes– CR116C and S6B– contained transposons within the *tgrBC* locus, but this was not consistent across other isolates. In addition, the boundaries of the unalignable regions differed in each genome (i.e. there were no regions used as repeated recombination hotspots). Instead, we conclude that the unalignable regions result from repeated, extensive genomic duplications, recombination events, and rearrangements within the *tgrBC* locus. This activity appears more intense in genomic regions closer to the paired, intact *tgrBC* genes.

Finally, we asked how these rearrangements shape the sequence and structure of *tgrBCs*, as matching *tgrB1* and *C1* protein sequences must heterodimerize for allorecognition in *D. discoideum* to occur(17, 21). We made alignments and phylogenies of the *tgrB* and *C* genes from the full-length, diverging gene pairs in each genome. Using GARD, we uncovered 12 sites of historical recombination within *tgrB1* genes and 10 sites in *tgrC1*; several of these recombination sites were found within the previously biochemically-defined regions of TgrB1 and TgrC1 that mediate binding and allorecognition(21) (Fig. 4E-F). These recombination sites separated regions of the *tgr* genes with different evolutionary histories. For example, a portion of the S6B TgrB1 had a *citrinum*-like N-terminal sequence and a *discoideum*-like C-terminal sequence (Fig. 4G). Additionally, in comparison to the whole genome tree, the *tgrC1* phylogeny had more well-supported discrepancies than the *tgrB1* tree did (Fig. S8). Thus, even though the strains analyzed here have low nucleotide divergence and high syntenic conservation genome-wide (Fig. 1 & 2), we see prolific gene gain, fragmentation, and loss of *tgrBCs*. We conclude that *Dictyostelium* generates the vast sequence diversity of its allorecognition genes via local genomic instability and prolific recombination.

## Discussion

To perform accurate self/non-self recognition, organisms require highly polymorphic cell surface factors, ideally those that continually generate new variants and specificities. Using our collection of high-quality, chromosome-scale genomes across diverse *Dictyostelium* species, we uncovered unusual features of *tgr* gene family evolution, particularly of *tgrBC* recognition genes. Some *tgrs* evolve slowly, indicating that these genes experience different evolutionary pressures and likely different molecular functions than *tgrB1* and *C1*. Other *tgr* subfamilies (*tgrAHIO* and *tgrBEJ/CDK*) are both large and evolutionarily dynamic, duplicating as often as the major *Dictyostelium* transposons (Fig. 3, Fig. S5). Moreover, we uncovered a role for intralocus recombination and local genomic rearrangements in creating this diversity. Complex arrays of new and partial *tgr* genes were repeatedly gained and lost, both from the *tgrBC* locus (Fig. 4) and the rest of the genome. Moreover, the local genomic instability in the *tgrBC* locus is long-lived and has continually shuffled the genomic content there for at least 50-100 million years. In addition, the intact *tgrBCs* are themselves conglomerates, resulting from recombination-mediated mixing and matching of different *tgr* genes and alleles. The recombination events occurred across the gene length, including within the biochemically-defined regions that mediate TgrB1/C1 binding and determine specificity(21).

Because polymorphic *tgrBs* and *tgrCs* bind each other as heterodimers, we initially assumed that new polymorphisms in *tgrB* would co-evolve with those in *tgrC*. However, it appears that these two proteins do not evolve in lockstep, despite their shared function in recognition. When we examined pairs of diverging *tgrBCs*, over 50% of these loci (13/25) had partial, truncated *tgrBs*, whereas nearly all (96%) had full length *tgrCs*. Even when we considered only gene pairs where both proteins were intact, *tgrB1s* had much less amino acid divergence than *tgrC1s* (Fig. S5), suggesting distinct selective pressures or functional constraints may be acting on each gene.

Prior evolutionary studies have analyzed sets of *Dictyostelium* ‘social genes’ together instead of studying *tgrs* individually, due to 1) the difficulty of disentangling different *tgr* orthologs with short read data, and 2) the low overall polymorphism rates among previously sequenced *D. discoideum* genomes, which necessitated analyzing gene pools. These studies have varied in their definitions of ‘social genes’ and methodology, leading to conflicting reports that *Dictyostelium* social genes evolve under relaxed selection (26), balancing selection (24), and/or rapid adaptive evolution (25). When analyzing sequence variation within the *tgr* genes, we did not find strong evidence of positive or balancing selection at the level of the full gene sequence. However, the complex recombination patterns we uncovered here are known to interfere with measures of selection, as each portion of a gene has a different evolutionary history and therefore a different expected rate of polymorphisms. This may account in part for the differing findings in previous studies. Analyses of more closely related *tgrBC* alleles may be useful for disentangling these selective signatures.

In addition, we propose that the structural variation itself is a critical part of *tgrBC* diversification. The *tgrBC* locus appears to be a highly active mutational hotspot, where repeated, local double-strand breaks (and the non-homologous repair of these breaks) generate rearrangements and large-scale gene gains and losses, both of *tgrBC*s and the neighboring genes. Yet, it is not clear how these double-strand breaks could be formed. In other species (e.g., mammalian antibody class switching(45), or yeast WHO elements(46)), such hotspots can result from the regulated binding of nucleases and repeated cuts made at a specific genomic locus. If a similar nuclease were binding within *tgrBC* and cleaving the adjacent region, that could explain both the long-lived nature of the hotspot across diverse species as well as its continued proximity to the intact, diverging *tgrBC* pairs. The mechanistic characterization of the generation and maintenance of this hotspot will be a fascinating topic for future studies.

How do new alleles for self/non-self recognition emerge? This question has been particularly mysterious for the *tgrB1/C1* heterodimer, as novel mutations in the recognition sequence of each locus would presumably cause mismatches with its cognate partner, leading to a failure of multicellular development. Our analyses suggest a potential solution to this problem and could explain why these recognition genes belong to such large gene families.

Prior work showed that mismatched, merodiploid *tgrB* and *tgrC* alleles can be expressed without disrupting allorecognition using other loci(17). Therefore, *tgr* genetic diversity could arise if a single, functional *tgrBC* pair enables multicellular development, while this locus continually generates full and partial *tgr* gene duplicates nearby. These adjacent duplicates would diverge rapidly (potentially as pseudogenes), accumulating mutations, deletions, and/or additional recombination events to produce novel variants. Occasionally, we speculate that fragments from these diverged *tgrs* could be recombined into a developmentally-expressed *tgrBC* locus, causing rapid sequence change. While this sort of mutation would presumably often generate non-matching, loss-of-function TgrB/C paired alleles, this would be consistent with the many partial *tgrB* and *C* loci we observed. Such regions could continue to evolve neutrally (as long as another *tgrBC* locus was still active during development) until the duplicates were either lost or compatible alleles arose in the partner gene, generating a new allorecognition module.

This proposal is in some ways similar to the hypothesis that social genes accumulate polymorphisms because they are rarely required and therefore experience relaxed selection(26). However, in our proposal, the *tgr* gene fragments could accumulate polymorphism neutrally, not only in the absence of cheater strains, but also when these gene fragments were physically distant from an intact *tgr* locus. If so, the locus would behave analogously to some bacterial toxin-immunity systems for self/non-self recognition, wherein unused, flanking genetic cassettes can recombine into a locus to rapidly alter toxin functions (47, 48). Many elements of animal polymorphic self/non-self recognition are also diversified by recombination, gene duplication, structural variation, such as the primate immunoglobulin, T-cell receptor, and MHC loci (49), the tunicate histocompatibility loci(50), and the cnidarian allorecognition genes(51). Our work uncovers an extreme example of this hyperdiversification in *Dictyostelium*, within a well-characterized model system for allorecognition, cell recognition, and cheater exclusion.

Finally, beyond the *tgr* genes, these genome assemblies and curated gene annotations offer a valuable resource for studying the evolution and function of all other gene families in *Dictyostelium*. These sequences are at evolutionary distances that enable the study of gene-level diversity and selection in Amoebozoa, allowing for analyses of selection and rapidly diversifying genes. We invite the community to make use of these resources for analyses of genome evolution, gene family diversification, and cell-cell recognition systems.

## Materials and Methods

Using a combination of Illumina reads and long Nanopore reads, we generated *de novo* genome assemblies and annotations for ten *Dictyostelium* isolates, which were obtained from the Dicty Stock Center (50). Nine genomes were for wild, group 4 *Dictyostelium* species, selected for their useful genetic distance to the *D. discoideum* AX4 reference genome(33). We also sequenced the axenic lab strain AX2-214 as well as full, closed genomes for two *Chlamydia*-like endosymbionts residing in the wild strains. Genome-scale synteny was analyzed using MCScanX(52). The *tgr* genes were classified based on MAFFT 7.471 alignments that were trimmed with TrimAL 1.2 before phylogeny construction using IQtree 1.6.12 (53–55). See SI Appendix, Extended Methods for full details on genome assembly and validation, alignments and phylogenetics, mating assays, and identification of *tgr* genes, BUSCO genes, orthology groups, centromeres, telomeres, mitochondrial, and rDNA.

## Data and software availability

Assembled genomes and annotations have been deposited in GenBank (SUB15159038 for *Dictyostelium* genomes, SUB13956370 for the *Chlamydia-*like symbiont in GS8B, SUB13956414 for the Chlamydia-like symbiont in M4B). All other data are available as supplemental files and custom scripts are available at: https://github.com/MayarMAhmed/dicty_genomes.

## Acknowledgments

We thank the members of the Levin lab, Gad Shaulsky, Chris Thompson, and Jason King for their suggestions on this project. M. Holland, J. Drurey, and T. Levin were supported by NIH R35-GM150681 and the Burroughs Wellcome Fund Investigators in the Pathogenesis of Infectious Disease award (T. Levin, P.I.). Janet Young was supported by NIH grant R01-GM74108 (Harmit S. Malik, P.I.).

## References

1. M. Paoletti, S. J. Saupe, C. Clavé, Genesis of a fungal non-self recognition repertoire. PLoS One 2, e283 (2007).

2. J. Zhao, et al., Identification of allorecognition loci in Neurospora crassa by genomics and evolutionary approaches. Mol. Biol. Evol. 32, 2417–2432 (2015).

3. M. H. Oberbarnscheidt, F. G. Lakkis, Innate allorecognition. Immunol. Rev. 258, 145–149 (2014).

4. R. D. Rosengarten, M. L. Nicotra, Model systems of invertebrate allorecognition. Curr. Biol. 21, R82–92 (2011).

5. P. Kundert, G. Shaulsky, Cellular allorecognition and its roles in Dictyostelium development and social evolution. Int. J. Dev. Biol. 63, 383–393 (2019).

6. A. Krensky, C. Clayberger, The nature of allorecognition. Curr. Opin. Nephrol. Hypertens. 2, 898–903 (1993).

7. B. Afzali, R. I. Lechler, M. P. Hernandez-Fuentes, Allorecognition and the alloresponse: clinical implications. Tissue Antigens 69, 545–556 (2007).

8. A. P. Gonçalves, et al., Allorecognition upon fungal cell-cell contact determines social cooperation and impacts the acquisition of multicellularity. Curr. Biol. 29, 3006–3017.e3 (2019).

9. A. P. Gonçalves, N. L. Glass, Fungal social barriers: to fuse, or not to fuse, that is the question. Commun. Integr. Biol. 13, 39–42 (2020).

10. D. Wall, Kin recognition in bacteria. Annu. Rev. Microbiol. 70, 143–160 (2016).

11. G. Glöckner, et al., The multicellularity genes of dictyostelid social amoebas. Nat. Commun. 7, 12085 (2016).

12. E. A. Ostrowski, M. Katoh, G. Shaulsky, D. C. Queller, J. E. Strassmann, Kin discrimination increases with genetic distance in a social amoeba. PLoS Biol. 6, e287 (2008).

13. R. Benabentos, et al., Polymorphic members of the lag gene family mediate kin discrimination in Dictyostelium. Curr. Biol. 19, 567–572 (2009).

14. J. L. Dynes, et al., LagC is required for cell-cell interactions that are essential for cell-type differentiation in Dictyostelium. Genes Dev. 8, 948–958 (1994).

15. Y. Wang, G. Shaulsky, TgrC1 has distinct functions in Dictyostelium development and allorecognition. PLoS One 10, e0124270 (2015).

16. S. Hirose, G. Chen, A. Kuspa, G. Shaulsky, The polymorphic proteins TgrB1 and TgrC1 function as a ligand-receptor pair in Dictyostelium allorecognition. J. Cell Sci. 130, 4002–4012 (2017).

17. S. Hirose, R. Benabentos, H.-I. Ho, A. Kuspa, G. Shaulsky, Self-Recognition in Social Amoebae Is Mediated by Allelic Pairs of Tiger Genes. Science 333, 467–470 (2011).

18. N. Gruenheit, et al., A polychromatic “greenbeard” locus determines patterns of cooperation in a social amoeba. Nat. Commun. 8, 14171 (2017).

19. M. Katoh-Kurasawa, P. Lehmann, G. Shaulsky, The greenbeard gene tgrB1 regulates altruism and cheating in Dictyostelium discoideum. Nat. Commun. 15, 3984 (2024).

20. H.-I. Ho, S. Hirose, A. Kuspa, G. Shaulsky, Kin recognition protects cooperators against cheaters. Curr. Biol. 23, 1590–1595 (2013).

21. G. Chen, et al., TgrC1 mediates cell-cell adhesion by interacting with TgrB1 via mutual IPT/TIG domains during development of Dictyostelium discoideum. Biochem. J. 452, 259–269 (2013).

22. R. Sucgang, et al., Comparative genomics of the social amoebae Dictyostelium discoideum and Dictyostelium purpureum. Genome Biol. 12, R20 (2011).

23. I. S. Hayakawa, K. Inouye, Species recognition in social amoebae. J. Biosci. 43, 1025–1036 (2018).

24. E. A. Ostrowski, et al., Genomic signatures of cooperation and conflict in the social amoeba. Curr. Biol. 25, 1661–1665 (2015).

25. S. Noh, K. S. Geist, X. Tian, J. E. Strassmann, D. C. Queller, Genetic signatures of microbial altruism and cheating in social amoebas in the wild. Proc. Natl. Acad. Sci. U. S. A. 115, 3096–3101 (2018).

26. J. L. de Oliveira, et al., Conditional expression explains molecular evolution of social genes in a microbe. Nat. Commun. 10, 3284 (2019).

27. Genome. NCBI. Available at: https://www.ncbi.nlm.nih.gov/datasets/genome/?taxon=5782. [Accessed 25 June 2025].

28. B. Edelbroek, J. Kjellin, J. Jerlström-Hultqvist, S. Koskiniemi, F. Söderbom, Chromosome-level genome assembly and annotation of the social amoeba Dictyostelium firmibasis. Sci. Data 11, 678 (2024).

29. G. Glöckner, et al., Sequence and analysis of chromosome 2 of Dictyostelium discoideum. Nature 418, 79–85 (2002).

30. R. M. McBee, S. A. Rozmiarek, N. R. Meyerson, P. A. Rowley, S. L. Sawyer, The effect of species representation on the detection of positive selection in primate gene data sets. Mol. Biol. Evol. 32, 1091–1096 (2015).

31. T. E. Douglas, M. R. Kronforst, D. C. Queller, J. E. Strassmann, Genetic diversity in the social amoeba Dictyostelium discoideum: population differentiation and cryptic species. Mol. Phylogenet. Evol. 60, 455–462 (2011).

32. F. Liu, et al., Novel symbionts reveal amoebae as significant hosts for environmental chlamydiae. bioRxiv 2024.11.01.621453 (2024).

33. L. Eichinger, et al., The genome of the social amoeba Dictyostelium discoideum. Nature 435, 43–57 (2005).

34. W. Makałowski, J. Zhang, M. S. Boguski, Comparative analysis of 1196 orthologous mouse and human full-length mRNA and protein sequences. Genome Res. 6, 846–857 (1996).

35. M. Manni, M. R. Berkeley, M. Seppey, E. M. Zdobnov, BUSCO: Assessing genomic data quality and beyond. Curr. Protoc. 1, e323 (2021).

36. I. M. Zada-Hames, Analysis of karyotype and ploidy of Dictyostelium discoideum using colchicine-induced metaphase arrest. J. Gen. Microbiol. 99, 201–208 (1977).

37. E. C. Cox, C. D. Vocke, S. Walter, K. Y. Gregg, E. S. Bain, Electrophoretic karyotype for Dictyostelium discoideum. Proc. Natl. Acad. Sci. U. S. A. 87, 8247–8251 (1990).

38. J. Cappello, K. Handelsman, H. F. Lodish, Sequence of Dictyostelium DIRS-1: an apparent retrotransposon with inverted terminal repeats and an internal circle junction sequence. Cell 43, 105–115 (1985).

39. K. Li, P. Xu, J. Wang, X. Yi, Y. Jiao, Identification of errors in draft genome assemblies at single-nucleotide resolution for quality assessment and improvement. Nat. Commun. 14, 6556 (2023).

40. R. D. Rosengarten, et al., Leaps and lulls in the developmental transcriptome of Dictyostelium discoideum. BMC Genomics 16, 294 (2015).

41. M. Stajdohar, et al., dictyExpress: a web-based platform for sequence data management and analytics in Dictyostelium and beyond. BMC Bioinformatics 18, 291 (2017).

42. K. L. Kindle, R. A. Firtel, Identification and analysis of Dictyostelium actin genes, a family of moderately repeated genes. Cell 15, 763–778 (1978).

43. J. Zucko, et al., Polyketide synthase genes and the natural products potential of Dictyostelium discoideum. Bioinformatics 23, 2543–2549 (2007).

44. G. Glöckner, et al., The complex repeats of Dictyostelium discoideum. Genome Res. 11, 585–594 (2001).

45. J. Stavnezer, C. E. Schrader, IgH chain class switch recombination: mechanism and regulation. J. Immunol. 193, 5370–5378 (2014).

46. M. Osborne, A. Fubara, E. Ó Cinnéide, A. Y. Coughlan, K. H. Wolfe, WHO elements - A new category of selfish genetic elements at the borderline between homing elements and transposable elements. Semin. Cell Dev. Biol. 163, 2–13 (2024).

47. J. Arenas, K. Schipper, P. van Ulsen, A. van der Ende, J. Tommassen, Domain exchange at the 3’ end of the gene encoding the fratricide meningococcal two-partner secretion protein A. BMC Genomics 14, 622 (2013).

48. S. J. Poole, et al., Identification of functional toxin/immunity genes linked to contact-dependent growth inhibition (CDI) and rearrangement hotspot (Rhs) systems. PLoS Genet. 7, e1002217 (2011).

49. D. Yoo, et al., Complete sequencing of ape genomes. Nature 1–18 (2025). 10.1038/s41586-025-08816-3.

50. A. W. De Tomaso, H. Rodriguez-Valbuena, Histocompatibility in Botryllus schlosseri and the origins of adaptive immunity. Immunogenetics 77, 22 (2025).

51. A. L. Huene, et al., A family of unusual immunoglobulin superfamily genes in an invertebrate histocompatibility complex. Proc. Natl. Acad. Sci. U. S. A. 119, e2207374119 (2022).

52. Y. Wang, et al., MCScanX: a toolkit for detection and evolutionary analysis of gene synteny and collinearity. Nucleic Acids Res. 40, e49 (2012).

53. K. Katoh, K. Misawa, K.-I. Kuma, T. Miyata, MAFFT: a novel method for rapid multiple sequence alignment based on fast Fourier transform. Nucleic Acids Res. 30, 3059–3066 (2002).

54. S. Capella-Gutiérrez, J. M. Silla-Martínez, T. Gabaldón, trimAl: a tool for automated alignment trimming in large-scale phylogenetic analyses. Bioinformatics 25, 1972–1973 (2009).

55. B. Q. Minh, et al., IQ-TREE 2: New models and efficient methods for phylogenetic inference in the genomic era. Mol. Biol. Evol. 37, 1530–1534 (2020).

