## Supporting Information for "Hypermutable hotspot enables the rapid evolution of self/non-self recognition genes in *Dictyostelium*"

### Extended Methods

#### *Dictyostelium* isolates sequenced

All *Dictyostelium* wild isolates were ordered from the Dicty Stock Center (1) and stored as frozen spore stocks in 10% DMSO (SIGMA-ALDRICH D2650-100ML). The wild isolates used in this study are KGL29A (DBS0305280), OT3A (DBS0304501), CR116C (DBS0304503), PJ11 (DBS0235829), AR5b (DBS0235745), CF3B (DBS0304504), GS8b (DBS0304535), M4B (DBS0304528), and S6B (DBS0304502). All wild strains were previously collected by Jim Cavender and John Landolt. We additionally used the axenic lab strain AX2-214 (DBS0235534).

#### Culturing and harvesting *Dictyostelium* cells

Wild *Dictyostelium* isolates are cultured on bacterial lawns of Gram-negative *Klebsiella pneumophila* (DBS0305928) spread on SM plates. To collect enough *Dictyostelium* cells for DNA extraction while limiting contamination by *Klebsiella*, we scraped the plates after *Dictyostelium* visually cleared the bacterial lawn but prior to fruiting body formation (as differentiated cells and spores are difficult to lyse for DNA extraction). For AR5b, CF3B, GS8b, M4B, and S6B: we additionally performed a centrifugation of the harvested cells at 100 x g for 1 min through 5 mL of  20% Percoll (Fisher Scientific 45-001-748) to physically separate the lighter bacterial cells from heavier *Dictyostelium* cells. Lab strain AX2-214 was grown axenically in HL5 (Formedium HLB0102) supplemented with penicillin and streptomycin (Fisher scientific 15-140-122).

#### Extracting *Dictyostelium* genomic DNA

For KGL29A, OT3A, CR116C, PJ11: harvested cells were resuspended in 10 mL of lysis buffer (100 mM NaCl, 50 mM Tris pH 7.5, 10 mM EDTA pH 8, 0.5% SDS, 0.3mg/mL proteinase K). The lysis was left rotating at room temperature for 1-2 days. Phenol/chloroform/isoamyl alcohol was added to the lysis buffer at a 1:2 ratio. The phenol/lysis mixture was inverted several times and then spun at 13,000 x g for 5 minutes. The supernatant above the phenol interface was collected (~4 mL), and isopropanol was added at a 1:1 ratio. The isopropanol/DNA mix was inverted several times to precipitate the gDNA. For KGL29A and OT3A, there was enough free-floating DNA to spool on glass hooks, which were then rinsed in 100% and 70% EtOH and left to air dry before resuspending in 1X TE. For CR116C and PJ11, the DNA was pelleted by spinning at 200g for 2 min. The pellet was rinsed with 100% and 70% EtOH. The pellet is then left to air dry overnight before resuspending in 1X TE. For all KGL29A, OT3A, CR116C, PJ11: once DNA was dissolved, 2 uL of 10 mg/mL RNAse was added for every 200 uL of TE/DNA and allowed to sit at RT overnight.

For AR5b, CF3B, GS8b, M4B, and S6B, and AX2-214: high molecular weight DNA was extracted using the Qiagen Genomic Tip 100/G columns (Qiagen 10243) following the yeast DNA extraction protocol with three modifications. First, we used $2x10^8$ *Dictyostelium* cells/mL in cell lysis C1 buffer to load the optimal amount of DNA on the Genomic Tip 100/G column. Second, we added 200 µg/ml RNaseA to the G1 digestion buffer prior to the 1 hour digestion. Lastly, the isopropanol precipitated DNA was washed in ice-cold 100% EtOH, then in 70% EtOH, and allowed to completely dry on a glass hook prior to resuspension in the final 1X TE solution. These modifications produced high-quality, high molecular weight DNA suitable for deep hybrid Nanopore and Illumina sequencing.

#### DNA sequencing and quality control

To sequence the gDNA, we ordered at least 1 Gbp of Oxford Nanopore long reads and 2 million reads of Illumina 150bp paired-end short reads (Table S1). To assess the quality of the sequencing reads, we ran fastqc(2) on the Illumina short reads and Nanoplot 1.0.0 (3, 4) for long reads. We filtered for high-quality Nanopore reads (quality score > 5 and length > 1000) with nanofilt 2.8.0(4). To filter for high-quality Illumina reads (Phred score > 30 and length > 140), we used Trimmomatic v0.38(5). *Dictyostelium* has a relatively low GC content (~25%) while *Klebsiella* bacteria generally have a GC content around 50%(6). To determine if a sample has high bacterial contamination, we examined the fastqc-generated distribution of read GC content for the presence of high %GC reads.

**Bacterial contamination filtration from sequencing and endosymbiont genome assembly**
Although we used multiple methods to physically separate bacteria from amoeba cells prior to the DNA extraction, the sequencing reads inevitably contained some bacterial contamination. To ensure the purity of the *Dictyostelium* reads used in our assemblies, we followed a decision tree to filter out as much bacterial contamination as possible. We first took the reads and generated an initial assembly using Flye v2.9(7) for a 34 Mbp genome. For AX2 and S6B, bacterial contamination was insignificant and the assembly was already high quality at this step. However, most strains still had significant bacterial contamination, and the initial assemblies were highly fragmented.

Therefore, to identify sources of bacterial contamination, we extracted the first 2 kbp of each contig and used blast+ BLASTN v2.7.1(8) with default parameters to search for matching bacterial genomes. The top-hit bacterial reference genome (typically strains of food bacteria such as *Klebsiella pneumoniae* or *Escherichia coli*) was downloaded from NCBI GenBank. The Nanopore reads were then mapped to these references to remove bacterial contaminant reads using minimap2 v0.2.4(9, 10). The remaining unmapped reads were used for *Dictyostelium de novo* genome assembly. For most strains, this was sufficient to remove the expected bacterial contamination inherent in our culturing methods and produce high quality *Dictyostelium* genomes.

However, for GS8b and M4B, this bacterial filtration step was unsuccessful and the sequences were heavily contaminated with unidentifiable bacterial DNA. We therefore performed an alternative filtration method to remove the unknown bacterial contamination. We first assembled a genome for the “food bacteria” strain using Flye (6 Mbp genome with scaffolds option) (7),(7) using the non-*Dictyostelium* reads from KGL29A, as the KGL29A sample had many high-quality Nanopore and Illumina reads. This produced a *Klebsiella pneumoniae* 5.2 Mbp genome with 2 plasmids (note: this strain is also called *Klebsiella aerogenes* by the Dicty stock center). We then mapped the Nanopore sequencing reads from GS8b and M4B with minimap2 to known organisms in the sample, *Dictyostelium (*reference AX4 genome (11)GCA_000004695.1) and *Klebsiella* food bacteria (our assembly). The remaining unmapped reads were still a mixture of *Dictyostelium* and unknown bacterial reads. Because almost half the unmapped sequencing reads were unknown bacterial contamination, we were able to assemble the remaining contaminant genome with Flye (6 Mbp genome setting). This produced a 1.5 Mbp *Chlamydia*-like genome for each of the GS8b and M4B samples, which was then used to filter out endosymbiont reads from the Nanopore long reads. To assemble the final *Dictyostelium* genome, we combined the AX4 mapped and remaining unmapped reads together and ran Flye with 34 Mbp settings. This method resulted in high-quality *Dictyostelium* and bacterial endosymbiont genomes for both GS8b and M4B. In PJ11, a *Chlamydia*-like genome was present but at lower read abundance, and so these additional assembly steps were not necessary to produce the *Dictyostelium* genome and a full *Chlamydia*-like genome was not recoverable.

To prepare the Illumina short reads for polishing the genome assemblies, we processed Illumina reads with the same contamination filtration steps as used for the Nanopore reads. To map Illumina reads to reference genomes, we used Burrows-Wheeler Aligner (bwa v0.7.17 and bwa-mem2 v2.2.1)(12, 13). Illumina short reads that were considered pure *Dictyostelium* were used in later polishing steps.

#### ***Dictyostelium*** **genome assembly and quality control**

For all isolates, filtered Nanopore reads were assembled with Flye using the 34 Mbp genome, scaffolds option. To assess the completeness of our genomes, we used BUSCO v5.3.2 (14, 15) to search for a collection of xxx conserved eukaryotic genes. All Flye assemblies achieved a BUSCO completeness score >90%. These assemblies were then iteratively polished ten times with filtered Illumina reads using pilon (16). Each polishing step was quality checked for completeness and k-mer error rate (to quantify any aberrant differences in k-mer abundances between the reads and assembled genomes) using BUSCO and k-mer abundance difference (KAD) (17) respectively. Assemblies with BUSCO completeness scores above 90% and fragmentation under 5% were considered high quality and complete, although the mitochondrial genome did not always assemble within a single contig (Table S3).

A small amount of bacterial contamination was still present in the filtered reads, resulting in many short bacterial contigs in the polished assemblies. We opted to only keep contigs that aligned to the reference AX4 genome when using MUMer v4.0 (18) with the default settings. Unaligned contigs were considered as contamination and excluded from further analysis and the Genbank submission pipeline. The genome statistics of the finalized assemblies were collected using Quast v5.0.2 (19, 20). An N50 >1 Mbp and an L50 <10 were considered high quality (compared to the reference AX4 *D. discoideum* genome [GCA_000004695.1] N50 of 5.5 Mbp and L50 of 3). To calculate the sequencing coverage, we mapped filtered Nanopore and Illumina reads to the final assembly using minimap2(9, 10) and BWA(12, 13), respectively. We then merged the alignments and calculated average read depth across the genome.

We expected all of the isolates to be haploid, similar to the AX4 reference strain. To check, we followed an all-sites variant calling approach using bcftool (21) (v1.2) in haploid mode with the illumina reads. We calculated per-site variant allele frequency for all positions to search for any read polymorphisms present at a frequency greater than 5%. In most genomes, we did not detect any variants that passed this frequency. The only exception was S6B, which had a small number of variants above this threshold, but none met the VCF criteria for variant calling. This result suggests that the sequenced isolates were indeed clonal and haploid.

#### ***Dictyostelium*** **gene prediction and annotation**

To predict gene sequences in each genome, we used GeneMarkEP+ (22) , which uses protein hints to improve *ab initio* gene finding in eukaryotic genomes. We provided the reference *D. discoideum* AX4 protein sequences (GCA_000004695.1) as the protein hint database. To identify rRNA genes, we used RNAmmer(23) with default parameters. After the gene prediction, we used MAKER(24–29) to generate a custom repeat library, which was provided to RepeatMasker(30) for softmasking each genome prior to annotation. Following the repeat masking step, we identified tRNA genes using tRNAscan-SE (31, 32). After the gene annotation was complete, we added functional annotation labels to each predicted coding sequence with results from BLASTP (8) of the UniProt database release 2024_01 (33) and InterProScan v5.57-90 (34, 35) of the PFAM database(36). Lastly, all annotation files were formatted with unique gene identifiers using scripts from the MAKER pipeline.

**Building the *Dictyostelium* whole genome tree**

To construct a phylogeny of all the sequenced genomes, we used the 174 single-copy BUSCO sequences of each species. We then aligned the coding sequences of each BUSCO ortholog across all genomes using MAFFT v7.471(37) with the auto-detect direction option, trimmed the alignments using gblocks v0.91b (38, 39) with default settings, and concatenated the ortholog alignments into an alignment matrix with phykit v1.19.3 (40). The best-fit model was identified with IQ-TREE2 (41) and used to construct the tree with 1000 bootstrap replicates and approximate likelihood-ratio test (aLRT) statistics. Additionally, we assessed the divergence of the dataset by calculating pairwise dS, pairwise nucleotide identity, and pairwise amino acid identity between each genome and *D. discoideum* AX4 using PAML CODEML v4.9h(42) on the alignment matrix of 174 orthologs that were present in every single species.

***Chlamydiae* endosymbiont classification and gene annotation**

To classify the endosymbiont genomes, we first identified their 16S sequences using BLASTN (8) to query our genomes with the 16S nucleotide sequence *of Chlamydia trachomati*s strain HAR-13 (NR_025888.1). The GS8b MH001 symbiont had one copy of the 16S rRNA sequence, while the M4B MH002 symbiont had three copies. The 16S sequence was not assembled for the PJ11 endosymbiont. Using the GS8b- and M4B-derived 16S sequences, we ran BLASTN on the NCBI web server with default settings to find the most closely related known bacterial species. From the BLASTN results, we took the top 10 non-redundant 16S sequences from each BLASTN search. All were part of the *Chlamydiae* phylum. To create a phylogenetic tree of representative *Chlamydiae*, we collected 42 *Chlamydiae* RefSeq representative 16S sequences and included one *Lentisphaerae* 16S as an outgroup (NR_146378.1). Using MAFFT v7.471(37), we performed an alignment of all the 16S sequences with autodetect direction and default setting. We ran IQ-TREE2 (41) to identify the best-fit model to construct the phylogenetic tree and generate support values with 1000 bootstrap and aLRT statistics. Lastly, we used Bakta v1.7.0 (43) with default settings for gene prediction and annotation of the bacterial endosymbiont genomes.

**Identifying mating types of wild *Dictyostelium* isolates**

The mating type locus of *Dictyostelium discoideum* is a single 5-6kbp region containing either *matA* (type 1), *matBCD* (type 2), or *matST* (type 3). In the AX4 reference genome, there is only a *matA* gene. We therefore used the mating type loci from other isolates of closely related *Dictyostelium discoideum* to provide reference genes of mating type 2 and 3 genes (*matBCD*: FN543121.1 and *matST*: FN543120.1). To define the mating types in our genomes, we first identified which ortholog groups contained the AX4 *matA* gene (DDB_G0289165), and the neighboring conserved genes of the mating type locus (DDB_G0289171 and DDB_G0289163/DDB_G0289161). Every isolate contained at least 1 homolog of the OG0000999 group and the locus neighboring genes OG0002508 and OG0000816. Our gene annotations number gene IDs in syntenic order so to collect all the genes in the locus, we collected the transcripts of all the genes with ID numbers between the two neighboring gene homologs. We then aligned all genes in the locus including the neighboring genes together with MAFFT 7.471(44) and generated a phylogenetic tree using FastTree 2.1.1(45). From the phylogenetic tree, clades of *matA-T* could be defined by clustering with the reference sequences of each gene. The mating types were identified by the presence of relevant homologs *matA* (type 1), *matCD* (type 2), *matST* (type 3). Some genomes also contained fusions of *matB* and *matD* or multiple *matT* genes, which may represent different mating types (Fig. S3).

#### Testing macrocyst formation in *Dictyostelium* wild isolate crosses

*Dictystelium* crosses were performed in SS buffer (0.6 g/L NaCl, 0.75 g/L KCl, 0.3 g/L CaCl2) on LP agar (1 g/L lactose, 1 g/L Bacto Peptone, 15 g/L agar powder) in a 24-well plate. Each *Dictyostelium* isolate was cultured by streaking spore stocks on bacterial lawns of Gram-negative *Klebsiella pneumophila* (DBS0305928) spread on SM plates. Once fruiting bodies formed, we collected a single fruiting body head (containing >1e6 spores) from each isolate for every cross. We then mixed the *Dictyostleium* spores for each cross at a 1:1 ratio and added them to a well with LP agar in triplicate. *Klebsiella pneumophila* was grown overnight in SM liquid culture and adjusted to an OD600 of 6 in the SS buffer. 5uL of concentrated *Klebsiella pneumophila* was then added to each well. The plates were wrapped in aluminum foil to prevent light exposure and incubated at 22C for 2 weeks. Macrocysts were imaged at 10x magnification after pipetting the supernatant of each well to break up any clumps.

#### Identification & validation of chromosomal rearrangements and duplications

To assess the structural similarities across genomes, we compared the synteny for each isolate vs. AX4 as well as the following pairs: AX4 with AX2-214, AX2-214 with CR116C, CR116C with OT3A, OT3A with M4B, M4B with S6B, S6B with CF3B, CF3B with KGL29A, KGL29A with GS8B, GS8B with PJ11, and PJ11 with *D. firmibasis* TNS-C-14. For each pair of isolates, we ran BLASTP to compare all predicted proteins between the two genomes (BLAST+ v2.10.1; e-value ≤ $1 \times 10^{-10}$; retaining the five highest-scoring hits per query). Additionally, we extracted the protein-coding gene coordinates from each genome's GFF3 annotation and reformatted them into a simple four-column table (contig, gene ID, start, and end).

We used MCScanX (v. 1.0) to chain BLASTP homology hits into collinear blocks defined by conserved gene order, enabling the systematic discovery of inversions, translocations, and tandem duplications(46). Analyses were run with default scoring parameters (MATCH_SCORE=50, MATCH_SIZE=5, GAP_PENALTY=0.1, and MAX_GAPS=25) and a block E-value threshold of $1\times10^{-4}$, providing high sensitivity for both genome-wide and locus-specific rearrangements. The resulting collinearity outputs were visualized using SynVisio, a web-based platform for exploring genome-wide synteny (47).

To validate these chromosomal rearrangements, we used the genome assembly quality validation tool CRAQ (Clipping-based Read Assembly Quality, v1.0.9) (48). CRAQ uses mapped sequencing reads to systematically identify regions likely to be misassembled versus true structural rearrangements by analyzing soft-clipped read clusters. For each genome, we mapped both our Illumina and Nanopore reads back to the assembly using BWA v0.7.15 (12) and Minimap2 v2.24 (10), respectively. Using these mapped sequences, we run CRAQ to identify the Clip-based Structural Errors (CSE; indicative of large-scale misassemblies) and Clip-based Regional Errors (CRE; indicative of small-scale misassemblies). Next, we examined the overlap between our Synteny-identified chromosomal rearrangements and the CRAQ-identified CSE regions to identify whether the rearrangements are from assembly artifacts or true rearrangements. CRAQ also calculated two Assembly Quality Indices (AQI) within non-overlapping 500kb windows per contig and average scores across the entire genome: Structural AQI (S-AQI), reflecting the structural integrity and presence of large-scale misjoins, and Regional AQI (R-AQI), indicating small-scale consensus accuracy. Both scores range from 0 to 100, with higher values indicating higher assembly quality. Assemblies with AQI scores above 90 are considered reference quality; scores between 80–90 are high quality; scores between 60–80 represent draft-quality assemblies; and scores below 60 suggest low-quality assemblies.

All assemblies exhibited robust read coverage (>99% of bases covered) and negligible low-confidence regions. Global Structural AQI (S-AQI) exceeded 90 for eight species (i.e, reference quality assembly) and was ~86 for OT3A and AR5B (i.e, high quality assembly) (Table S2). In contrast, Regional AQI (R-AQI) widely varied, ranging from near zero in S6B to ~89 in GS8B. Two assemblies (GS8B, KGL29A) achieved high-quality R-AQI (80–90), three (CF3B, AX2-214, PJ11) were draft quality (60–80), and the remaining five (CR116C, OT3A, M4B, S6B, AR5B) scored below 60. These lower R-AQI values reflect pervasive small-scale consensus errors in some genomes that did not compromise overall scaffold structure. Across all genomes, CRAQ flagged 24 structural error breakpoints (CSEs): AR5B contained six; OT3A, five; CF3B, CR116C, and S6B, three each; and AX2-214, KGL29A, M4B, and PJ11, one each. Crucially, none of the synteny-defined rearrangement breakpoints overlapped CRAQ-flagged CSEs, indicating that the detected inversions and translocations are unlikely to be misassembly artifacts. Moreover, when we checked for overlaps between the CRAQ-flagged CSE regions and the tgrBC locus, none of the CSEs were located within the marker genes that defined the locus. The AQI scores for each assembly are summarized in Table S2.

**Identifying transposons, ribosomal regions, and mitochondria**
To comprehensively characterize the distribution of transposable elements (TEs) within our newly assembled *Dictyostelium* genomes, we employed a homology-based strategy using an Amoebozoa-specific TE library(49). Edelbroek et al. derived this library from Repbase, a comprehensive database of repetitive elements in eukaryotes (50), specifically extracting TE sequences from *Dictyostelium discoideum*, *D. firmibasis*, and related Amoebozoan species. These extracted sequences were filtered to retain only entries greater than 500 bp in length, ensuring enrichment for full-length, protein-coding TEs.

Using the library (TE_amoebozoa_500_ddis.fa) as a query, we performed TBLASTX searches against each genome assembly. Specifically, we applied stringent search parameters (e-value cutoff of $1\times10^{-15}$) to maximize identification accuracy and limit false positives. Among identified TEs, we specifically prioritized DIRS1 retrotransposons, as these elements frequently cluster at centromeric regions in *Dictyostelium* and would thus provide essential landmarks to assess chromosome structure and assembly completeness (51, 52). To systematically define DIRS1 clusters, we aggregated filtered TBLASTX hits into genome-wide sliding windows, each 10 kb in size, incremented by 5 kb steps. Windows containing ≥50 DIRS1 hits were flagged as high-density clusters, indicative of potential centromeric regions. Finally, we cross-referenced these high-density DIRS1 windows with orthologous gene predictions generated using OrthoFinder for validation.

To determine whether our newly assembled genomes contained the same telomeric repeats previously identified in the *D. firmibasis (49)* (5’-GAGGAGAGAGTCCCTTTTTTT-3’ and 5’-GGGGAGAGACAGGGGAGAGACA-3’), we first identified all tandem repeats and their consensus sequences using the Tandem Repeats Finder v4.09.1 (TRF) (53) with default parameters. Next, we used short-query BLASTN (8) v2.13.0 to look for the previously reported telomeric repeat sequences against the TRF-derived consensus sequences. To maximize detection sensitivity for degenerate or slightly imperfect telomere arrays, we relaxed mismatch penalties (-penalty -1) and gap costs (-gapopen 2), raised the e-value cutoff to 1000, and retained only hits with alignment identity ≥ 70 %. We parsed those BLAST hits to keep only exact coordinate matches within 15 kb of either end of a contig—defining them as candidate telomeric regions.

To determine if we had assembled the mitochondrial and/or rDNA loci, we used homology-based similarity searches guided by the mitochondrial (54) and rDNA (55) sequences from the *D. discoideum* AX4 reference genome, obtained from dictyBase (56, 57). For the mitochondrial loci, we searched our assemblies using two complementary approaches. First, we performed BLASTN v2.13.0 searches against the reference mitochondrial genome to detect contigs with significant nucleotide-level homology (e-value ≤ $1{\times}10^{-10}$). Second, we performed TBLASTN (8) v2.13.0 using the complete set of predicted mitochondrial proteins (e-value ≤ $1{\times}10^{-5}$), allowing us to identify contigs that encoded homologs of known mitochondrial proteins. We then intersected the two results to identify contigs supported by both DNA and protein evidence, increasing confidence that these contigs represent authentic mitochondrial sequences. For rDNA loci, we performed BLASTN(8) (v2.13.0) searches (e-value ≤ $1{\times}10^{-10}$) with the spliced rRNA transcript sequences, enabling us to identify contigs containing 17S, 5.8S, 26S, and 5S rRNA genes. The contigs corresponding to mitochondrial and rDNA loci are summarized in Table S3.

**Identifying and phylogenetically classifying *tgr* homologs**
To identify intact *tgr* homologs across the genomes, we used TBLASTN to search a blast+ custom database containing the predicted genes from all 13 genomes from Ax4 to *D. purpureum*, using the amino acid sequences from the 38 AX4 Tgr homologs as queries with an e-value cut-off of 1E-20. This resulted in 777 unique hits. After extracting the predicted transcripts for these 777 hits, we examined the sequences to determine if the coding sequence was in frame. For nine *D. purpureum* transcripts, we trimmed off the 5' and 3' UTRs to obtain in-frame sequences. Eleven other sequences were partial and out of frame. We added one or two Ns to the start of these sequences to put them in frame, then translated the dataset to obtain amino acid sequences.

To determine whether our use of AX4 reference *tgr* sequences limited our ability to recover hits in the more distantly related species, we then took our collection of *D. purpureum tgr* sequences and used them as TBLASTN queries in the same database. This search recovered the 777 sequences already found, plus 15 additional matches. The additional 15 transcripts were all within *D. purpureum*, and were short (<200 AA), with e-values close to our statistical cutoffs. We therefore concluded that the *D. purpureum* search did not result in additional full-length *tgr* homologs. See Table S4 for the gene IDs and classifications of all *tgr* homologs.

We aligned the set of 777 amino acid sequences using MAFFT 7.471 (37) and trimmed the resulting alignment with TrimAL 1.2 (58) using a gap threshold of 0.2. We constructed initial phylogenetic trees to explore the dataset using FastTree 2.1.1 (45). We also used hmmscan (24) to predict protein domains encoded by the sequences using the Pfam-A HMM database and an e-value threshold of 1E-3. When we looked at the 777 sequence alignment, we saw that some of the hits did not align well and/or included an aligned region that was considerably shorter than the 600-1200AA typical of most Tgr genes. To eliminate these presumably partial and/or pseudogene hits from our dataset, we created subsets of these sequences and iteratively aligned, trimmed, and made new trees until only a set of 500 well-aligned sequences remained. Through this process, we removed sequences with <300 aligned amino acids, any sequences with an alignment length of <650, and long branches that aligned poorly and did not contain the TIG repeats typical of Tgr proteins. We also identified 30 instances wherein two adjacent gene models encoded the N and C-terminus of a full-length *tgr* gene. We removed these 'split' *tgr* gene models from the dataset.

Using the curated set of 500 amino acid sequences, we generated an alignment with MAFFT, trimming alignment with TrimAL as above. We then constructed a phylogeny using IQtree 1.6.12 (41, 59) using the following parameters: -m TEST -bb 1000 -safe -nt AUTO. The best fit model was WAG+F+G4 (60). We used the consensus tree from IQtree to classify *tgr* genes into families. To do so, we identified well-supported subclades of the phylogeny and looked for the presence of AX4 *tgr* genes. We named each subclade according to the AX4 homologs found within it (e.g. if the clade contained *tgrA1, A2, A3, A4,* and *A5*, we named it the *tgrA* clade). If the subclade contained no AX4 homologs, we named it according to its nearest AX4 clades (e.g. tgrBEJ outgroup).

**Classifying *Dictyostelium* orthology groups and running diversity analyses**

To call orthology groups in our set of 12 genomes, we ran an all vs. all comparison with OrthoFinder 3.0 (61) using default settings. See Table S5 and S6 for gene ID categorization into orthology groups. For the 15 largest orthology groups, we did a preliminary classification of the gene family based on the AX4 homologs' classification in dictybase(56, 57) and domain architecture based on PFAM domain content.

To assess amino acid divergences within the *tgr* families and across other orthology groups, we first aligned protein sequences of each *tgr* gene family. We then exported a pairwise identity matrix using Geneious Prime version 2025.0. To pare down the *tgrB* and *tgrC* genes to only the likely functional alleles, we identified gene pairs where adjacent, diverging *tgrBEJ* genes were on the opposite strand from *tgrCDK* and both genes were approximately full length relative to AX4 *tgrB1/C1*. Most genomes contained one such pair and *D. intermedium* PJ11 encoded two pairs. We compared these genes to the single-copy *tgr* groups at the nucleotide level. We first aligned the amino acid sequence of each single-copy orthogroup with MAFFT v7.526(44) with default settings and converted to codon-aware nucleotide alignments with pal2nal v14 (62). These alignments were trimmed using trimal v1.5.rev0 (58) with a gap threshold of 0.2. Egglib v3.3.4 (63) was used to calculate pi, thetaW and Tajima's D for all ortholog groups. Any sequences with fewer than 100 analysed sites were discarded, while ortholog groups with less than 3.5 average utilised sequences were ignored in the Tajima's D calculations. To calculate dN/dS, we used PAML CODEML v4.10.7(42) model 0 to estimate a single dN/dS value for each trimmed ortholog group alignment. See Fig S5 and Tables S7-S9 for relevant statistics.

**Synteny, alignments, and recombination within *tgrBC* locus**

To identify the *tgrBC* locus across genomes, we first defined flanking marker genes that were consistently present and syntenic on either side of the *tgrBC* locus: *wacA* (*DDB_G0280537*) on the 5' end, and *chdB* (*DDB_G0280705*) and/or *DDB_G0280717* on the 3' end. We then used MCScanX and Synvisio to conduct a synteny analysis of the *tgrBC* locus. After identifying the three conserved flanking genes in each assembly, we extracted a genomic window centered on the locus, with 100 kb upstream and 100 kb downstream of the flanking markers, using a custom Python script (tgr_extraction.py), yielding locus-specific FASTA and GFF3 files. These segments underwent the same phylogeny-guided BLASTP and MCScanX pipeline (E-value ≤ $1\times10^{-4}$; default scoring) as described above. Visualization in SynVisio enabled the resolution of fine-scale structural variations, including tandem duplications, inversions, and gene-conversion events, at the tgrBC locus.

While this analysis gave an automated, coarse view of the duplications and rearrangements, it was still unclear how the *tgrBC* locus dynamics altered the local gene repertoires. We therefore selected pairs of closely related genomes to create alignments of the region. Because of the many differences in gene presence/absence as well as duplications, inversions, and rearrangements, typical alignment software was not able to create alignments of these >100 kb regions. Therefore, we used Mauve(64), which can better analyze large-scale recombination and gene gain/loss events. We extracted the region between the marker genes from each genome, then aligned these two regions using the progressive Mauve algorithm in Geneious Prime with these parameters: automatically calculate seed weight, automatically calculate minimum LCB score, assume collinear genomes, full alignment(64). To test for recombination signatures within the intact, paired *tgrBC* homologs, we aligned the transcripts for each gene with MAFFT and analyzed the alignment with GARD(65).

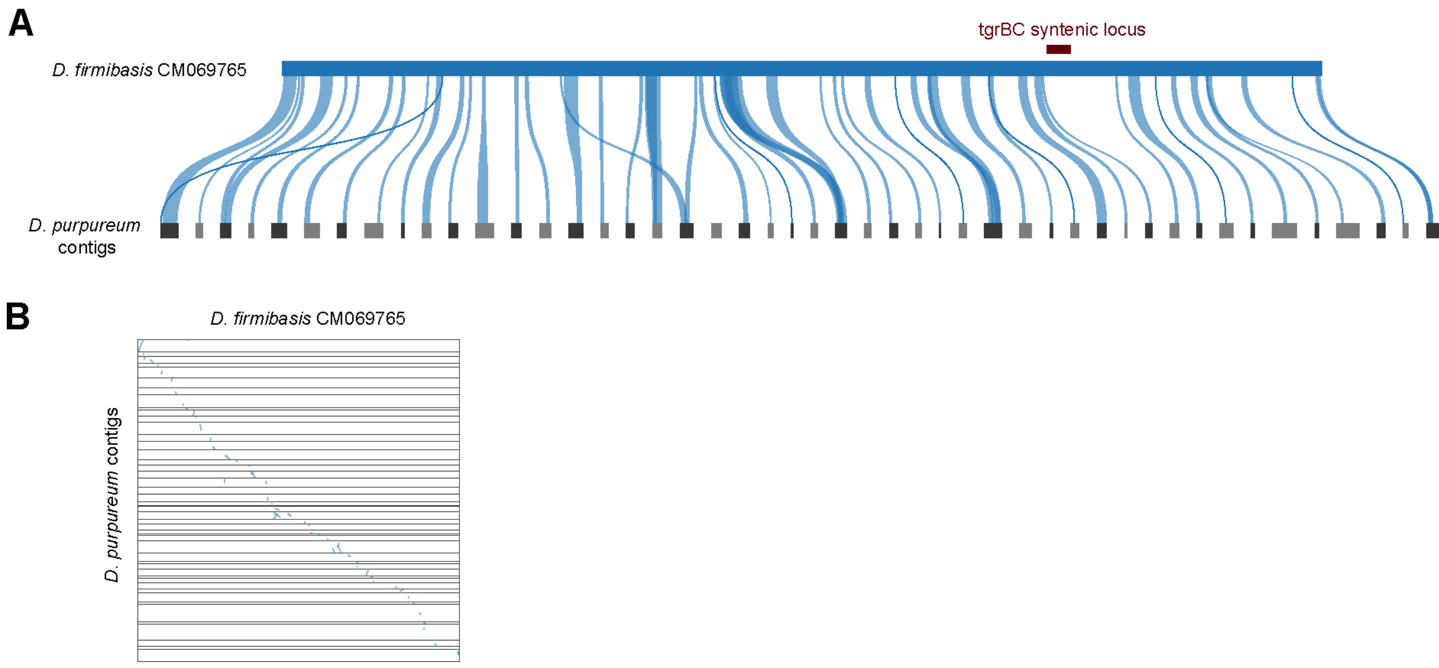


**Figure S1**. The *Dictyostelium purpureum* genome assembly is highly fragmented and incomplete, with over 85% of predicted genes only present as partial fragments, leading to difficulty identifying homologs or syntenic blocks. **A)** Regions of synteny (blue connecting lines) between *D. firmibasis* chromosome 1(Genbank assembly: GCA_036169595.1) and largest corresponding contigs from the current *D. purpureum* genome assembly (GCA_000190715.1). None of the syntenic regions overlap with the *tgrBC* syntenic locus **B)** Synteny visualized by dotplot using the blastp, MCScanX, Synvisio pipeline. Horizontal lines indicate the boundaries between *D. purpureum* contigs.

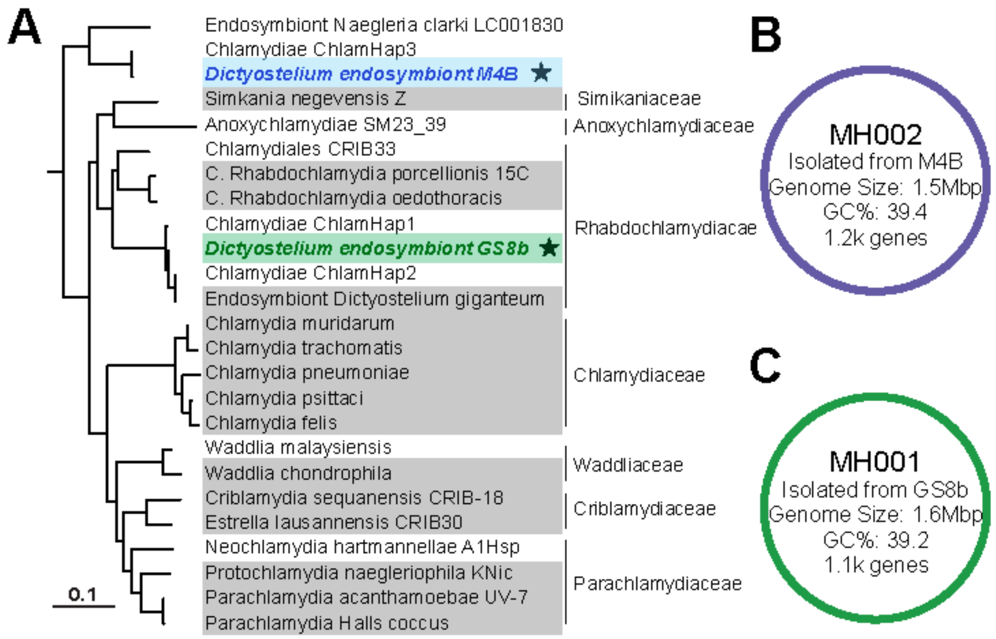


**Figure S2**. ***Chlamydia*-like endosymbiont genomes recovered from *Dictyostelium* amoebae. A)** 16S phylogenetic tree of *Chlamydia*-like endosymbionts. Symbiont genomes were selected to represent the major chlamydial families defined in Dharamshi et al 2023. Full genomes assembled prior to this study are highlighted in grey while all other samples were partial genomes or only 16S sequences. Symbiont genomes from this study are colored, bolded, and starred. The 16S sequence of the PJ11-associated symbiont was not recoverable from the partial genome and not included. **B-C)** Genome statistics of the full circular endosymbiont genomes isolated from *Dictyostelium* isolates M4B (**B**) and GS8b (**C**)

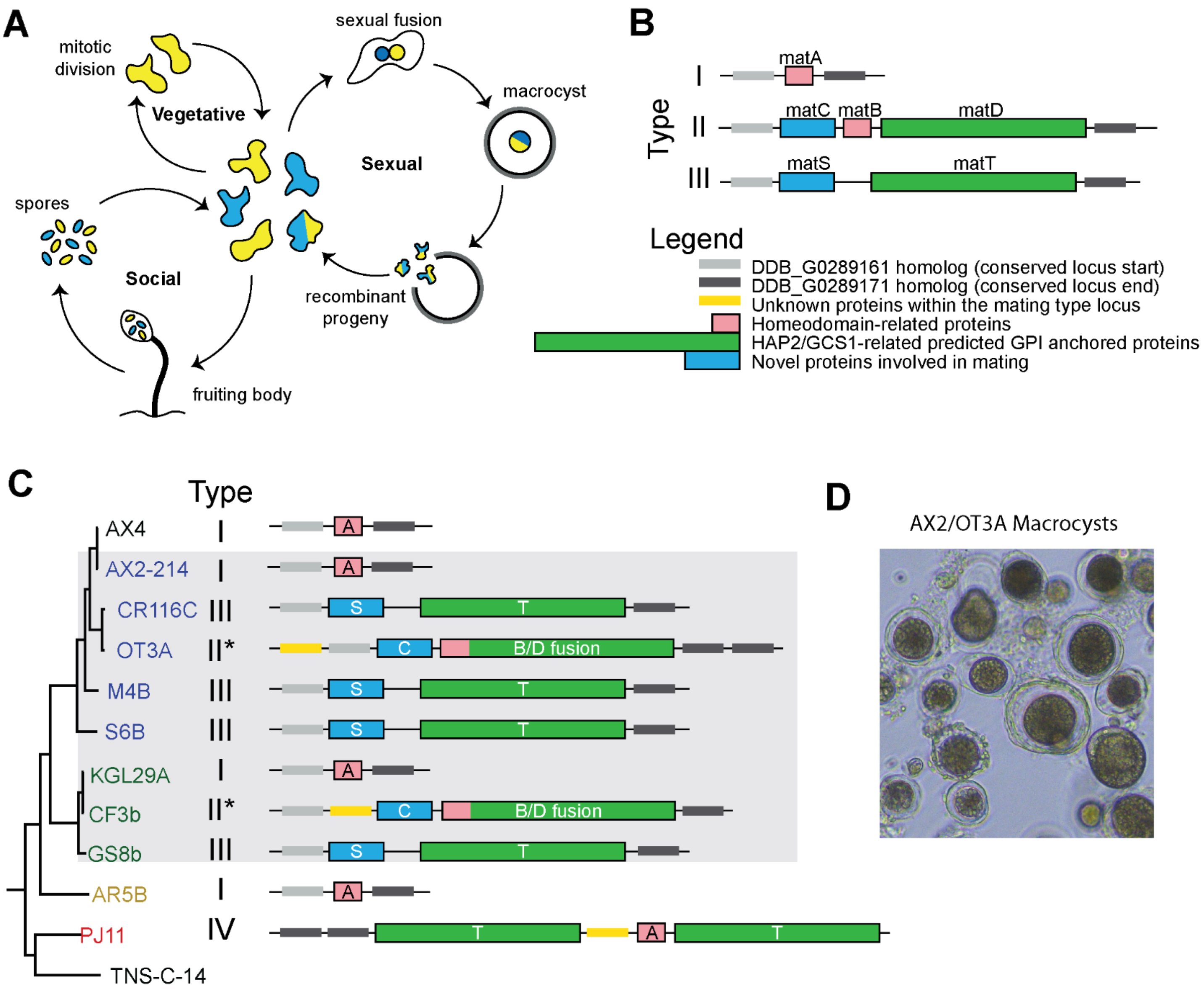


| | *discoideum* clade | | | | | *citrinum* clade | | |
|---|---|---|---|---|---|---|---|---|
| | I | II* | III | III | III | I | II* | III |
| | AX2 | OT3A | CR116C | M4B | S6B | KGL29A | CF3B | GS8B |
| AX2 | | ++ | + | - | - | | - | |
| OT3A | | | - | - | - | - | | |
| CR116C | | | | | | | - | |
| M4B | | | | | | | | |
| S6B | | | | | | | | |
| KGL29A | | | | | | | + | + |
| CF3B | | | | | | | | - |
| GS8B | | | | | | | | |

**Figure S3. Sexual life cycle and crosses of discoideum and citrinum clade *Dictyostelium spp*.** A) Schematic of the three different life cycles of *Dictyostelium discoideum*: 1) vegetative, asexual growth, 2) social multicellular development of *tgrBC* compatible strains, and 3) sexual fusion of *mat* mating type compatible strains. Blue and yellow colors indicate different genotypes, which recombine only during the sexual cycle. B) *Dictyostelium discoideum* has three described mating types, each with a different gene complement in a conserved mat locus. C) Mating type locus across the genomes assembled in this study, including some that had a fusion between the *matB* and *matD* genes labeled II*. The PJ11 *D. intermedium* genome had an unusual locus, which we call Type IV. D) Representative macrocyst images from the AX2/OT3A cross E) Summary of macrocyst formation across all crosses, with >10 macrocysts per well (++), 2-10 macrocysts (+), or no macrocysts observed(-). Dark gray boxes indicate pairs where crosses were not attempted. The citrinum clades crossed with each other and AX2 crossed with OT3A and CR116C, while M4B and S6B formed few to no macrocysts in any pair. However, we found that the type III strains generally mated with less efficiency than type I/II pairs (e.g. for CR116C and GS8B), so this may contribute to low mating efficiency in M4B and S6B.

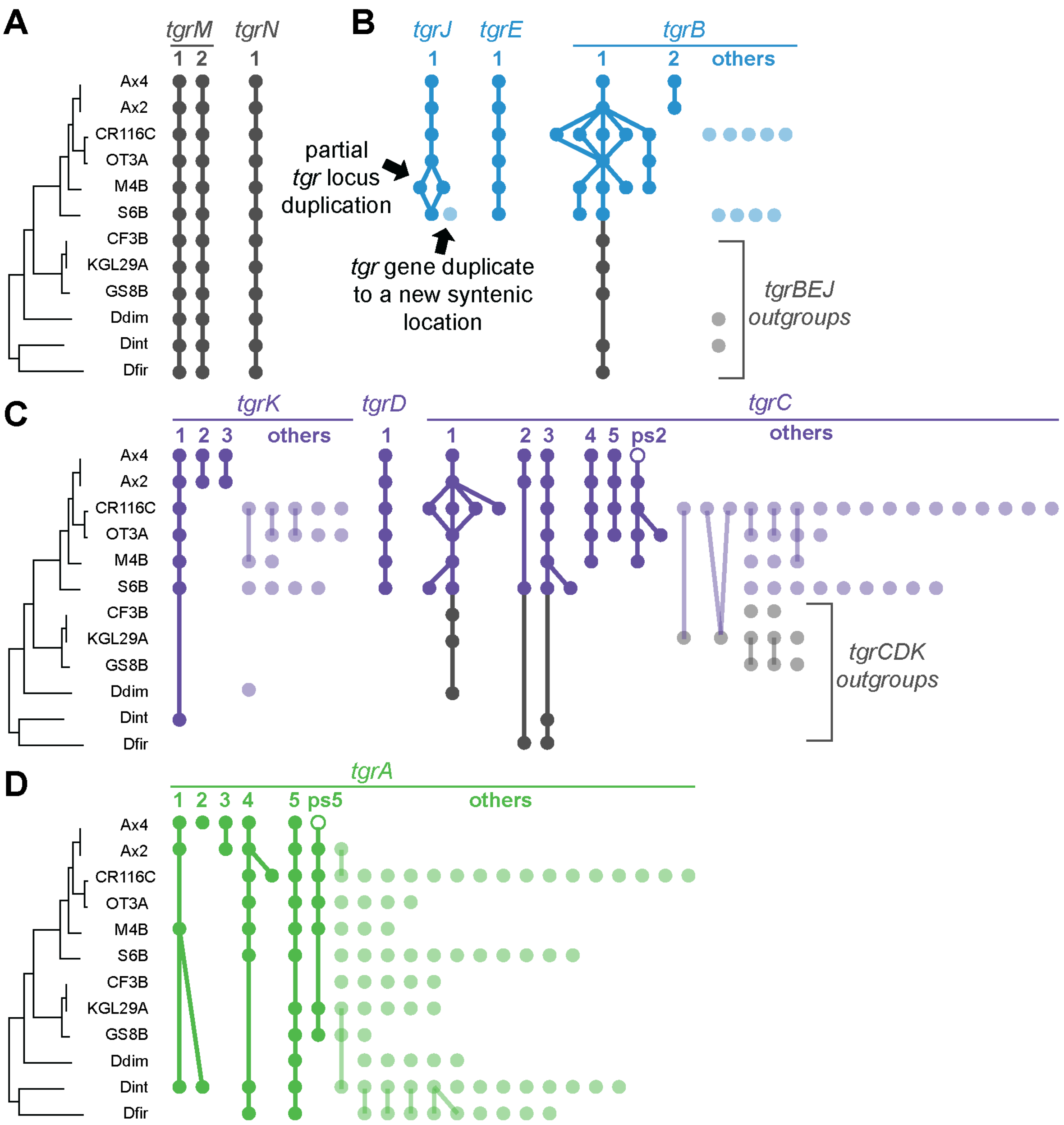


**Figure S4. Diverse patterns of duplication and retention in different *tgr* gene families**. For each genome, gene presence is indicated with a dot, and dots are connected when the homologs are present in the same syntenic location. Labels at the top of each column indicate the AX4 gene name (e.g. "*tgrM1*" for the first column of A). Lighter colored dots represent homologs that do not share synteny with AX4 *tgr* genes, according to MCScanX analyses. Open circles are those called as pseudogenes in AX4. **A)** Non-dynamic genes such as *tgrN* and *M* are single-copy and syntenic across all genomes. **B-C)** In contrast, *tgrB* and *tgrC* genes are frequently duplicated. Branched lines indicate genomes where multiple regions of one genome are syntenic with one region of another genome, due to repeated, partial duplications of the *tgr* locus. The *tgrDE* genes were conserved and unchanging, whereas *tgrJKs* exhibited some additional duplications. **D)** The *tgrA* family has undergone many gene duplications, generating a very large family with often lineage-specific chromosomal locations.

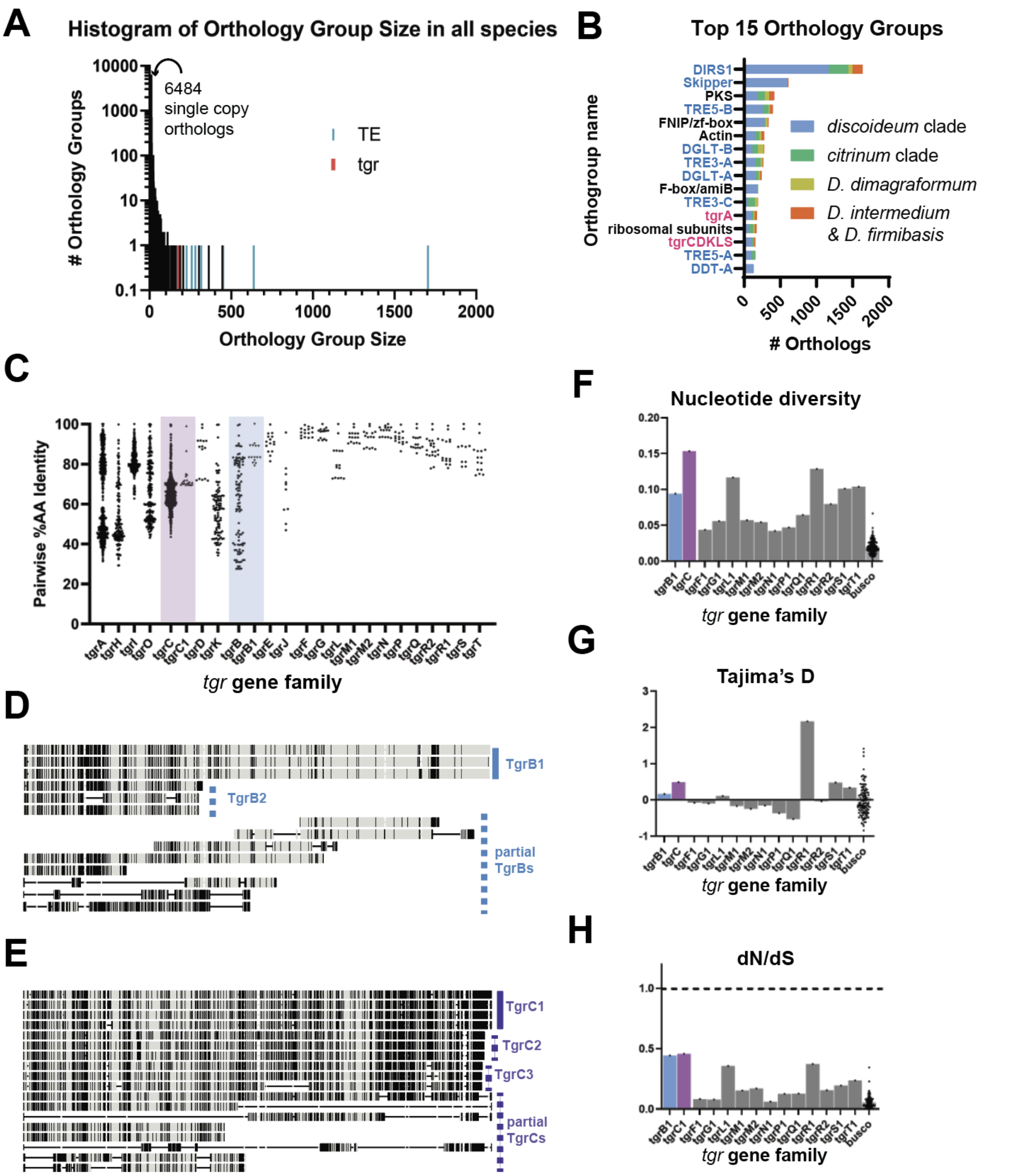


**Figure S5**. **Orthogroup composition and *tgr* gene diversity analyses**. **A)** Histogram of orthogroup size distribution across the 12 *Dictyostelium* genomes. In the 15 largest orthogroups, transposable elements (TEs) are shown in blue, *tgr* genes in red, and other protein-coding genes in black. **B)** Gene count in the 15 largest orthogroups, including transposable elements (blue text), *tgrA* and *tgrCDKLS* (pink text), and several protein-coding gene families such as polyketide synthases (PKS) and actin. The distribution of each gene family across different clades is indicated by bar colors: *discoideum* clade (blue), *citrinum* clade (green), *D. dimigraformum* AR5B (yellow), and *D. intermedium* PJ11 + *D. firmibasis* (orange). **C)** All-by-all pairwise amino acid identities within *tgr* gene families in the *discoideum* clade. The distributions for *tgrB* and *tgrC* were wide because they included numerous truncated duplicates (see **D** and **E**). Therefore, we also analyzed the paired, diverging *tgrB1* and *tgrC1* genes separately (see *tgrB1* and *tgrC1* columns). **D-E)** Amino acid alignments of the TgrB (**D**) and Tgr C (**E**) loci found in the *tgrBC* locus of AX4, OT3A, and CR116C. Regions of agreement with a consensus sequence are in light gray and regions of disagreement are indicated by black lines. TgrB1 and C1 are full length. TgrB2, C2, and C3 are shorter homologs named in AX4, while additional partial TgrB and C sequences are only in OT3A and CR116C. **F-H**) Nucleotide diversity and selection analyses with *discoideum* clade sequences: (**F**) nucleotide diversity (π/lseff), (**G**) Tajima's D neutrality test, and (**H**) dN/dS selection analysis. To avoid analyzing partial gene fragments or pseudogenes, only the paired, diverging *tgrB1* and *tgrC1* genes were included in these analyses (blue and purple bars) as compared to the other single-copy *tgr* genes and conserved BUSCO genes for reference.

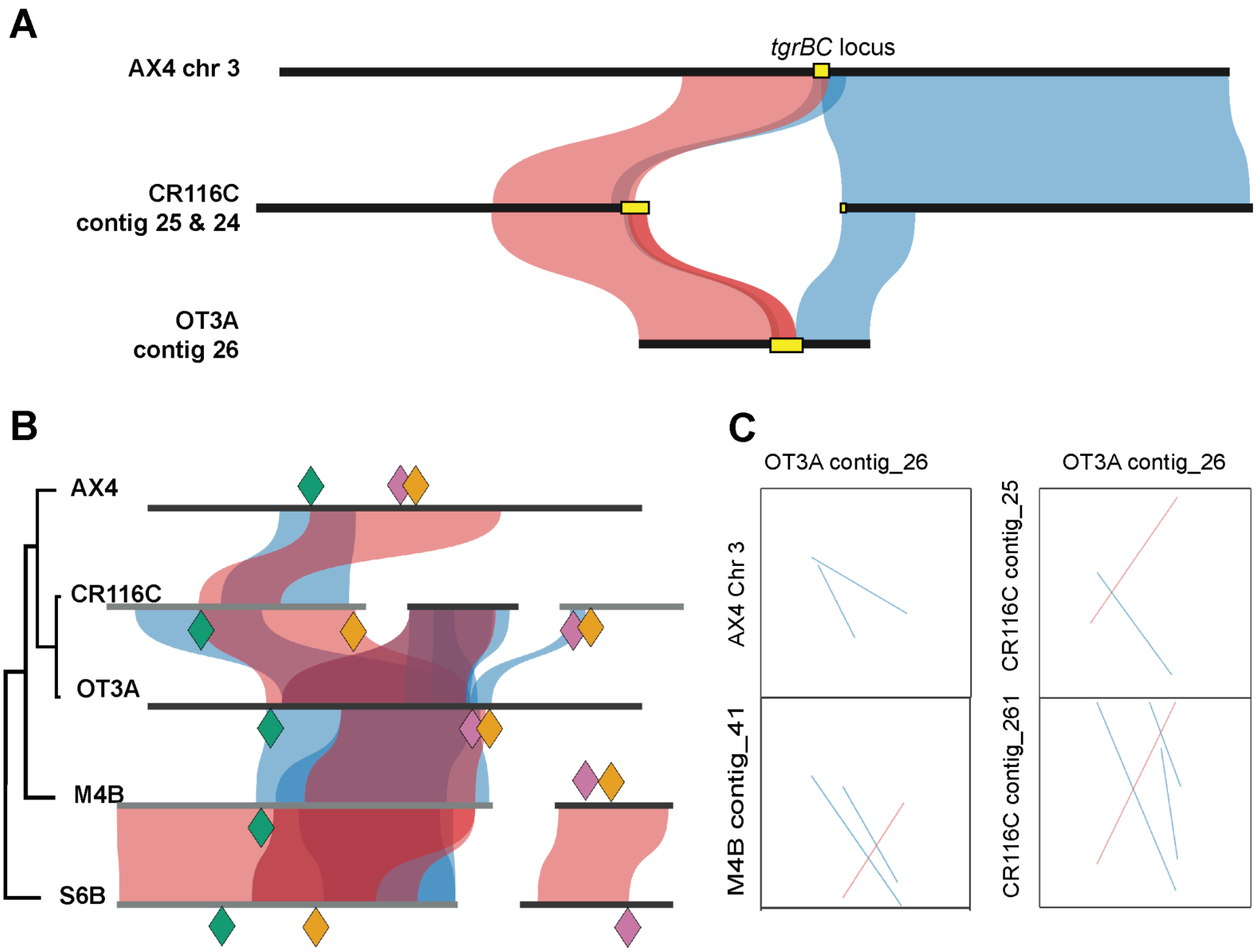


**Figure S6. Local, complex synteny within the *tgrBC* locus**. **A)** Chromosome/contig-wide synteny determined by MCScanX and SynVisio. Within *D. discoideum,* AX4 chromosome 3 (6.3Mb in length) is largely syntenic with corresponding regions of CR116C and OT3A. Blue, curved ribbons connect synteny matches in the same (forward) orientation as the reference, while red ribbons mark inverted synteny. However, this synteny becomes more complex, with multiple overlapping forward and reverse synteny predictions at the *tgrBC* locus (yellow box). **B)** In an attempt to disentangle the *tgrBC* locus, we examined the synteny using MCScanX at a more zoomed-in scale. Each horizontal bar depicts the tgrBC locus plus 100 kb of flanking sequence on either side. Colored diamonds show the locations of three flanking marker genes: wacA (green) at the 5' end and chdB (purple) and DDB_G0280717 (orange) at the 3' end. In CR116C, M4B, and S6B, the contig assembly breaks in the middle of the *tgrBC* locus. **C–D)** Representative dot plots for selected pairwise comparisons illustrate overlapping inversions and duplications within the locus. Axes list the compared contigs; blue lines denote forward matches and red lines denote inverted matches. When lines cross, this means that synteny matches are being called between the regions on both the forward and reverse strands simultaneously.

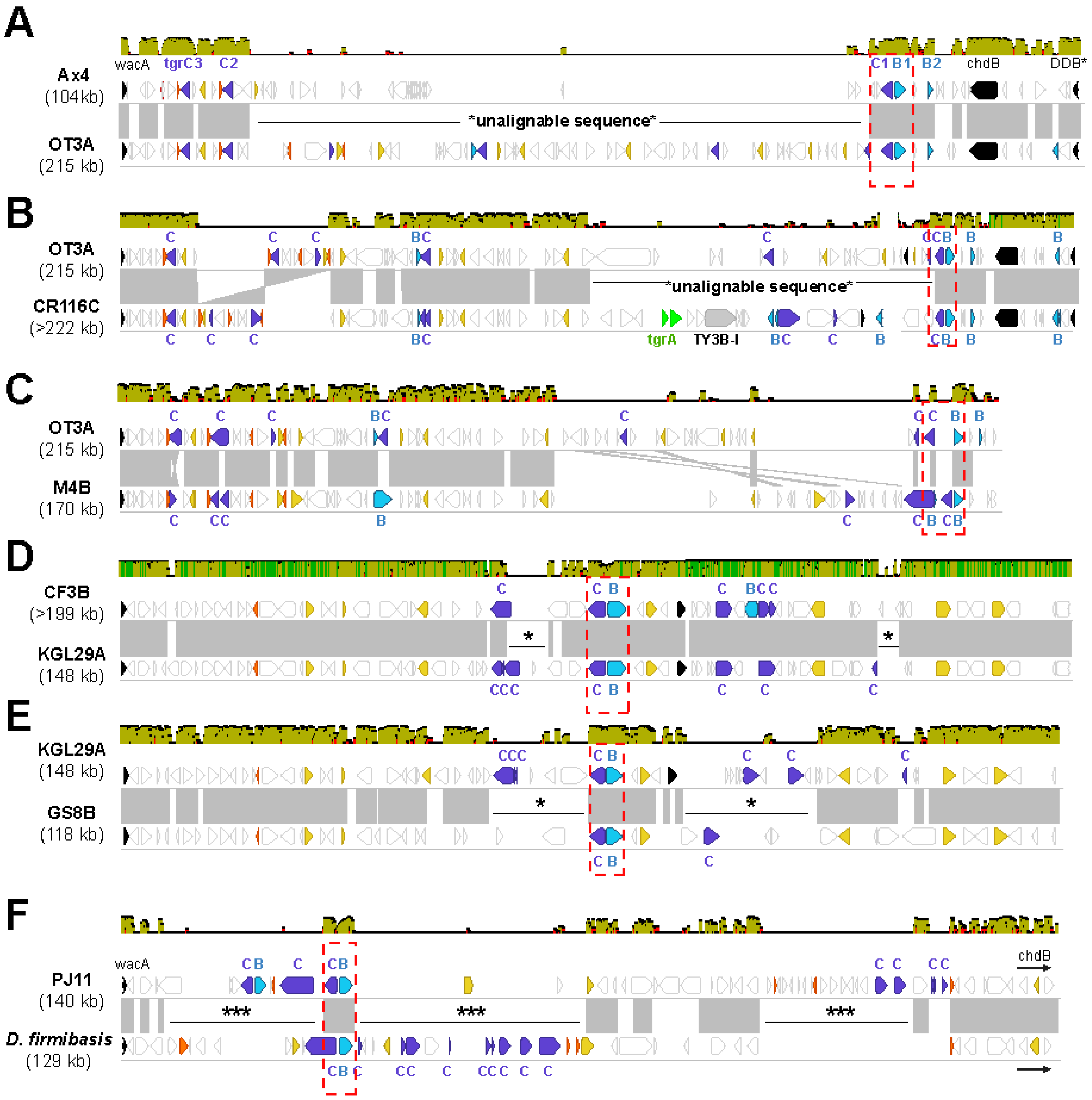


**Figure S7**. **Pairwise *tgrBC* alignments among additional strains**. All genes in the locus are block arrows colored as in Fig 5: *tgrB* = blue, *tgrC* = purple, marker genes = black, neighboring *arr* duplicates = orange, *gta* duplicates = yellow. Paired intact *tgrBCs* are indicated with a red dashed box. Grey blocks indicate regions of 30% nucleotide identity or higher, with inversions as in B shown as hourglass shapes. The percent nucleotide identity is also shown with the colored graph above each pairwise alignment: green = regions of 100% identity, yellow = 30-99% identity, red = <30% identity, and white showing unaligned regions with gaps. In CR116C, two *tgrA* alleles and a predicted transposon TY3B-I enter the locus in the unalignable sequence. In other species, the unalignable regions contain many different genes, often including multiple copies or fragments of *tgrC*.

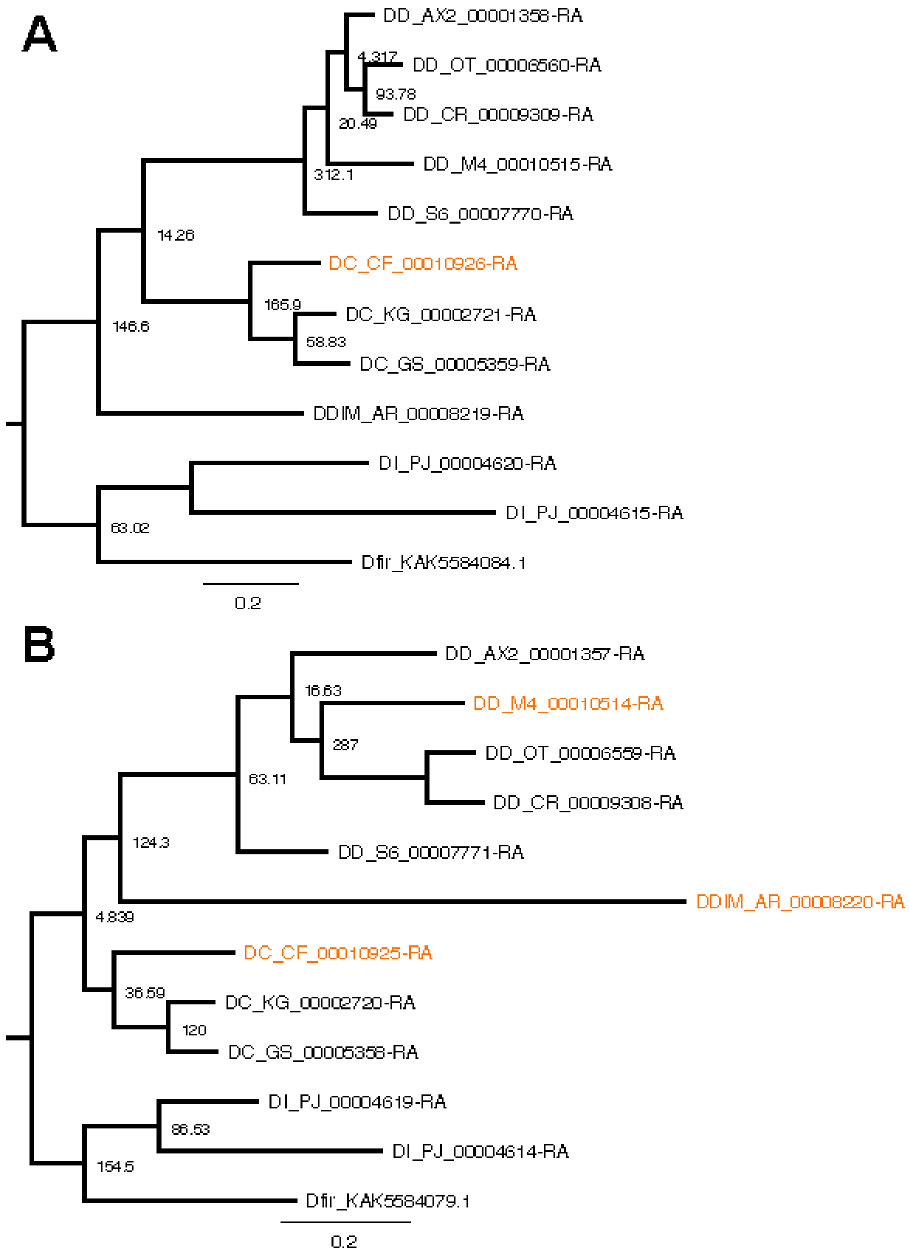


**Figure S8**. **Maximum likelihood phylogenetic trees of paired, diverging *tgrBs* (A) and *tgrCs* (B) show minor discordances with the species trees**. Tips are labeled with gene IDs, indicating the genome of origin (e.g. DD for *discoideum* clade sequences, DC for *citrinum* clade sequences). Branches that are discordant with the species tree in Fig 1 are indicated in orange. Nodes are labeled with aLRT statistics from PhyML.

### SI References


1. P. Fey, R. J. Dodson, S. Basu, R. L. Chisholm, One stop shop for everything Dictyostelium: dictyBase and the Dicty Stock Center in 2012. *Methods Mol. Biol.* **983**, 59–92 (2013).

2. S. Andrews, FastQC: A Quality Control Tool for High Throughput Sequence Data [Online]. (2010). Available at: http://www.bioinformatics.babraham.ac.uk/projects/fastqc/ [Accessed 23 June 2025].

3. W. De Coster, S. D'Hert, D. T. Schultz, M. Cruts, C. Van Broeckhoven, NanoPack: visualizing and processing long-read sequencing data. *Bioinformatics* **34**, 2666–2669 (2018).

4. W. De Coster, R. Rademakers, NanoPack2: population-scale evaluation of long-read sequencing data. *Bioinformatics* **39** (2023).

5. A. M. Bolger, M. Lohse, B. Usadel, Trimmomatic: a flexible trimmer for Illumina sequence data. *Bioinformatics* **30**, 2114–2120 (2014).

6. C. A. Ouellette, R. H. Burris, P. W. Wilson, Deoxyribonucleic acid base composition of species of Klebsiella, Azotobacter and Bacillus. *Antonie Van Leeuwenhoek* **35**, 275–286 (1969).

7. M. Kolmogorov, J. Yuan, Y. Lin, P. A. Pevzner, Assembly of long, error-prone reads using repeat graphs. *Nat. Biotechnol.* **37**, 540–546 (2019).

8. C. Camacho, *et al.*, BLAST+: architecture and applications. *BMC Bioinformatics* **10**, 421 (2009).

9. H. Li, New strategies to improve minimap2 alignment accuracy. *Bioinformatics* **37**, 4572–4574 (2021).

10. H. Li, Minimap2: pairwise alignment for nucleotide sequences. *Bioinformatics* **34**, 3094–3100 (2018).

11. L. Eichinger, *et al.*, The genome of the social amoeba Dictyostelium discoideum. *Nature* **435**, 43–57 (2005).

12. H. Li, R. Durbin, Fast and accurate short read alignment with Burrows-Wheeler transform. *Bioinformatics* **25**, 1754–1760 (2009).

13. N. Alganmi, H. Abusamra, Evaluation of an optimized germline exomes pipeline using BWA-MEM2 and Dragen-GATK tools. *PLoS One* **18**, e0288371 (2023).

14. M. Manni, M. R. Berkeley, M. Seppey, F. A. Simão, E. M. Zdobnov, BUSCO update: Novel and streamlined workflows along with broader and deeper phylogenetic coverage for scoring of eukaryotic, prokaryotic, and viral genomes. *Mol. Biol. Evol.* **38**, 4647–4654 (2021).

15. M. Manni, M. R. Berkeley, M. Seppey, E. M. Zdobnov, BUSCO: Assessing genomic data quality and beyond. *Curr. Protoc.* **1**, e323 (2021).

16. B. J. Walker, *et al.*, Pilon: an integrated tool for comprehensive microbial variant detection and genome assembly improvement. *PLoS One* **9**, e112963 (2014).

17. C. He, *et al.*, Factorial estimating assembly base errors using k-mer abundance difference (KAD) between short reads and genome assembled sequences. *NAR Genom. Bioinform.* **2**, lqaa075 (2020).

18. G. Marçais, *et al.*, MUMmer4: A fast and versatile genome alignment system. *PLoS Comput. Biol.* **14**, e1005944 (2018).

19. A. Mikheenko, A. Prjibelski, V. Saveliev, D. Antipov, A. Gurevich, Versatile genome assembly evaluation with QUAST-LG. *Bioinformatics* **34**, i142–i150 (2018).

20. A. Mikheenko, V. Saveliev, P. Hirsch, A. Gurevich, WebQUAST: online evaluation of genome assemblies. *Nucleic Acids Res.* **51**, W601–W606 (2023).

21. P. Danecek, *et al.*, Twelve years of SAMtools and BCFtools. *Gigascience* **10** (2021).

22. T. Brůna, A. Lomsadze, M. Borodovsky, GeneMark-EP+: eukaryotic gene prediction with self-training in the space of genes and proteins. *NAR Genom. Bioinform.* **2**, lqaa026 (2020).

23. K. Lagesen, *et al.*, RNammer: consistent annotation rRNA genes genomic sequences. *Nucleic Acids Res* (2007).

24. S. R. Eddy, Accelerated profile HMM searches. *PLoS Comput. Biol.* **7**, e1002195 (2011).

25. HMMER. Available at: http://hmmer.org/ [Accessed 23 June 2025].

26. C. Holt, M. Yandell, MAKER2: an annotation pipeline and genome-database management tool for second-generation genome projects. *BMC Bioinformatics* **12**, 491 (2011).

27. J. M. Flynn, *et al.*, RepeatModeler2 for automated genomic discovery of transposable element families. *Proc. Natl. Acad. Sci. U. S. A.* **117**, 9451–9457 (2020).

28. M. S. Campbell, C. Holt, B. Moore, M. Yandell, Genome annotation and curation using MAKER and MAKER-P. *Curr. Protoc. Bioinformatics* **48**, 4.11.1–4.11.39 (2014).

29. B. L. Cantarel, *et al.*, MAKER: an easy-to-use annotation pipeline designed for emerging model organism genomes. *Genome Res.* **18**, 188–196 (2008).

30. R. Hubley, *RepeatMasker: RepeatMasker is a program that screens DNA sequences for interspersed repeats and low complexity DNA sequences* (Github).

31. P. P. Chan, B. Y. Lin, A. J. Mak, T. M. Lowe, tRNAscan-SE 2.0: improved detection and functional classification of transfer RNA genes. *Nucleic Acids Res.* **49**, 9077–9096 (2021).

32. T. M. Lowe, S. R. Eddy, tRNAscan-SE: a program for improved detection of transfer RNA genes in genomic sequence. *Nucleic Acids Res.* **25**, 955–964 (1997).

33. UniProt Consortium, UniProt: The universal protein knowledgebase in 2025. *Nucleic Acids Res.* **53**, D609–D617 (2025).

34. P. Jones, *et al.*, InterProScan 5: genome-scale protein function classification. *Bioinformatics* **30**, 1236–1240 (2014).

35. M. Blum, *et al.*, InterPro: the protein sequence classification resource in 2025. *Nucleic Acids Res.* **53**, D444–D456 (2025).

36. J. Pei, *et al.*, Bridging the gap between sequence and structure classifications of proteins with AlphaFold Models. *J. Mol. Biol.* **436**, 168764 (2024).

37. K. Katoh, K. Misawa, K.-I. Kuma, T. Miyata, MAFFT: a novel method for rapid multiple sequence alignment based on fast Fourier transform. *Nucleic Acids Res.* **30**, 3059–3066 (2002).

38. J. Castresana, Selection of conserved blocks from multiple alignments for their use in phylogenetic analysis. *Mol. Biol. Evol.* **17**, 540–552 (2000).

39. G. Talavera, J. Castresana, Improvement of phylogenies after removing divergent and ambiguously aligned blocks from protein sequence alignments. *Syst. Biol.* **56**, 564–577 (2007).

40. J. L. Steenwyk, *et al.*, PhyKIT: a broadly applicable UNIX shell toolkit for processing and analyzing phylogenomic data. *Bioinformatics* **37**, 2325–2331 (2021).

41. B. Q. Minh, *et al.*, IQ-TREE 2: New models and efficient methods for phylogenetic inference in the genomic era. *Mol. Biol. Evol.* **37**, 1530–1534 (2020).

42. Z. Yang, PAML 4: phylogenetic analysis by maximum likelihood. *Mol. Biol. Evol.* **24**, 1586–1591 (2007).

43. O. Schwengers, *et al.*, Bakta: rapid and standardized annotation of bacterial genomes via alignment-free sequence identification. *Microb. Genom.* **7** (2021).

44. K. Katoh, D. M. Standley, MAFFT multiple sequence alignment software version 7: improvements in performance and usability. *Mol. Biol. Evol.* **30**, 772–780 (2013).

45. M. N. Price, P. S. Dehal, A. P. Arkin, FastTree 2--approximately maximum-likelihood trees for large alignments. *PLoS One* **5**, e9490 (2010).

46. Y. Wang, *et al.*, MCScanX: a toolkit for detection and evolutionary analysis of gene synteny and collinearity. *Nucleic Acids Res.* **40**, e49 (2012).

47. V. Bandi, C. Gutwin, Interactive Exploration of Genomic Conservation. *Proc. Graph. Interface* 74–83 (2020).

48. K. Li, P. Xu, J. Wang, X. Yi, Y. Jiao, Identification of errors in draft genome assemblies at single-nucleotide resolution for quality assessment and improvement. *Nat. Commun.* **14**, 6556 (2023).

49. B. Edelbroek, J. Kjellin, J. Jerlström-Hultqvist, S. Koskiniemi, F. Söderbom, Chromosome-level genome assembly and annotation of the social amoeba Dictyostelium firmibasis. *Sci. Data* **11**, 678 (2024).

50. W. Bao, K. K. Kojima, O. Kohany, Repbase Update, a database of repetitive elements in eukaryotic genomes. *Mob. DNA* **6**, 11 (2015).

51. J. Cappello, K. Handelsman, H. F. Lodish, Sequence of Dictyostelium DIRS-1: an apparent retrotransposon with inverted terminal repeats and an internal circle junction sequence. *Cell* **43**, 105–115 (1985).

52. G. Glöckner, *et al.*, The complex repeats of Dictyostelium discoideum. *Genome Res.* **11**, 585–594 (2001).

53. G. Benson, Tandem repeats finder: a program to analyze DNA sequences. *Nucleic Acids Res.* **27**, 573–580 (1999).

54. S. Ogawa, *et al.*, The mitochondrial DNA of Dictyostelium discoideum: complete sequence, gene content and genome organization. *Mol. Gen. Genet.* **263**, 514–519 (2000).

55. R. Sucgang, *et al.*, Sequence and structure of the extrachromosomal palindrome encoding the ribosomal RNA genes in Dictyostelium. *Nucleic Acids Res.* **31**, 2361–2368 (2003).

56. S. Basu, P. Fey, D. Jimenez-Morales, R. J. Dodson, R. L. Chisholm, dictyBase 2015: Expanding data and annotations in a new software environment: total dictybase upgrade 2015. *Genesis* **53**, 523–534 (2015).

57. P. Fey, R. J. Dodson, S. Basu, E. C. Hartline, R. L. Chisholm, dictyBase and the Dicty Stock Center (version 2.0) - a progress report. *Int. J. Dev. Biol.* **63**, 563–572 (2019).

58. S. Capella-Gutiérrez, J. M. Silla-Martínez, T. Gabaldón, trimAl: a tool for automated

alignment trimming in large-scale phylogenetic analyses. *Bioinformatics* **25**, 1972–1973 (2009).

59. D. T. Hoang, O. Chernomor, A. von Haeseler, B. Q. Minh, L. S. Vinh, UFBoot2: Improving the ultrafast bootstrap approximation. *Mol. Biol. Evol.* **35**, 518–522 (2018).

60. S. Kalyaanamoorthy, B. Q. Minh, T. K. F. Wong, A. von Haeseler, L. S. Jermiin, ModelFinder: fast model selection for accurate phylogenetic estimates. *Nat. Methods* **14**, 587–589 (2017).

61. D. M. Emms, S. Kelly, OrthoFinder: phylogenetic orthology inference for comparative genomics. *Genome Biol.* **20**, 238 (2019).

62. M. Suyama, D. Torrents, P. Bork, PAL2NAL: robust conversion of protein sequence alignments into the corresponding codon alignments. *Nucleic Acids Res.* **34**, W609–12 (2006).

63. M. Siol, T. Coudoux, S. Ravel, S. De Mita, EggLib 3: A python package for population genetics and genomics. *Mol. Ecol. Resour.* **22**, 3176–3187 (2022).

64. A. E. Darling, B. Mau, N. T. Perna, Progressive Mauve: Multiple alignment of genomes with gene flux and rearrangement. *arXiv [q-bio.GN]* (2009).

65. S. L. Kosakovsky Pond, D. Posada, M. B. Gravenor, C. H. Woelk, S. D. W. Frost, GARD: a genetic algorithm for recombination detection. *Bioinformatics* **22**, 3096–3098 (2006).
